# Genome-wide rules of transcription factor cooperativity revealed through *in silico* binding site ablation

**DOI:** 10.1101/2025.06.19.660093

**Authors:** Xuening He, Hjörleifur Einarsson, Petra Páleníková, Sarah Rennie, Dewei Hu, Ruining Cui, Christian Vaagensø, Kasper Lage, Robert Krautz, Robin Andersson

**Affiliations:** Section for Computational and RNA Biology, Department of Biology, University of Copenhagen, Denmark; The Novo Nordisk Foundation Center for Genomic Mechanisms of Disease, Broad Institute of MIT and Harvard, USA; Novo Nordisk Foundation Center for Protein Research, University of Copenhagen, Denmark; Stanley Center for Psychiatric Research, Broad Institute of MIT and Harvard, USA; Institute of Biological Psychiatry, Mental Health Services, Copenhagen University Hospital, Denmark

## Abstract

Transcription factor (TF) cooperativity plays a critical role in gene regulation. However, the underlying genomic rules remain unclear, calling for scalable methods to characterize the TF binding site (motif) syntax of regulatory elements. Here, we introduce DeepCompARE, a lightweight model paired with an *in silico* ablation (ISA) framework for genome-wide analysis of regulatory sequences. Our framework enables precise interpretation of the motif syntax governing chromatin accessibility, enhancer activity, and promoter function. We find that most TF motifs are pairwise independent, indicating a default additive behavior of TFs, and define a cooperativity score to quantify deviations from this baseline. This reveals synergy and redundancy as opposite effects along the same cooperative spectrum. TF redundancy is linked to promoter activity and broad expression, whereas TF synergy is associated with enhancer activity, physical interactions, and cell-type specificity. Our framework provides a quantitative model for TF cooperativity, offering new insights into gene regulatory logic.

## Introduction

Gene regulatory elements are essential for controlling the timing, specificity, and level of gene expression^1–4^. This control is encoded in regulatory DNA through specific arrangements of binding site sequences (motifs) for transcription factors (TFs) that regulate chromatin accessibility and the recruitment and processivity of the transcriptional machinery^5,6^. The resulting TF motif syntax is thought to influence the activity of gene regulatory elements by facilitating cooperativity between TFs^7–13^. Characterizing the motif syntax of gene regulatory elements and understanding how it influences TF cooperativity is therefore important for understanding transcriptional control.

Cooperativity between two or more TFs occurs when there is a functional dependency or coordination between them, i.e., when their combined functional activity is different from the aggregated effects independently asserted by each partner. For instance, co-binding of Forkhead box (FOX) and Myocyte Enhancer Factor 2 (MEF2) TFs boosts enhancer activity in lymphoblastoid cells^13^. TF cooperativity can also yield specific activities to gene regulatory elements only when certain conditions are met^13,14^. For example, the interferon β (IFN-β) enhancer requires co-binding of Activating Transcription Factor

2 (ATF2), Interferon Regulatory Factors 3 and 7 (IRF3, IRF7), c-Jun, and Nuclear Factor kappa-light-chain-enhancer of Activated B cells (NF-kβ) for activation, and lacking binding of any one entirely abrogates *IFN-β* gene activity^9^. Similarly, ETS Variant TF 2 (Etv2) and FoxC2 drive endothelial-specific gene expression through cooperative activation of enhancers^15^. Such encoded synergy between TFs can thus ensure the temporal, spatial, and cell type specificity of regulatory activity.

Gene regulatory elements can also encode TF redundancy, in which binding of either of two TFs is sufficient to exert full regulatory activity. As their combined functional activity differs from the sum of their independent effects, this is also a form of cooperativity. However, compared to synergy, redundancy only clearly manifests itself when the binding or availability of one TF is impaired and is compensated by the other. This way, TF redundancy may confer robustness to gene regulation in the face of variation in DNA sequence, cellular context, and environmental stress^16–18^. For example, housekeeping promoters are associated with low gene expression variability and often harbor multiple sites for redundant binding of Erythroblast Transformation Specific (ETS) family TFs^19,20^. In addition, redundant binding of Universal Stripe Factors maintains chromatin accessibility of regulatory elements even under adverse conditions^21^.

Recent advances in deep learning have enabled accurate modeling of regulatory activity from DNA sequences and accelerated discovery of motif cooperativity by providing functional readouts for any *in silico* DNA perturbation. Two main approaches to resolve cooperativity have been established: bottom-up approaches embed motifs into background sequences to measure the gain of regulatory activity upon introducing a new motif^22–24^, whereas top-down approaches often use natural genomic sequences and measure the loss of regulatory activity upon perturbation of an existing motif^25,26^. However, the syntactic constraints of TF binding sites and the quantitative effects of TF cooperativity at regulatory elements beyond a few TF pairs remain unknown, and a systematic investigation of genome-wide rules of TF cooperativity is still lacking.

The challenge is at least fourfold. First, regulatory activity is better predicted by nonlinear deep learning models than linear models^27^, indicating substantial nonlinearity between TF pairs. Yet, a baseline model to describe the default behavior of TF pairs is lacking. Second, predicting the effects of genome-wide perturbations of all TF combinations requires immense computation. Third, the trade-off between model complexity and prediction accuracy makes it hard to develop an accurate yet small deep learning model using conventional methods. Fourth, definitions of TF cooperativity and independence are diverse and can be contradictory between studies^25,26,28^, calling for uniform and formal definitions. The subtle yet critical difference between bottom-up and top-down approaches to infer cooperativity complicates the issue (Methods).

To address these challenges, we here develop a lightweight convolutional neural network for deep comparative analysis of regulatory elements (DeepCompARE), and a novel interpretation strategy, *in silico* ablation (ISA), for systematic analysis of the DNA sequence determinants of chromatin accessibility, enhancer activity, and promoter activity through computational masking of single nucleotides or motifs (**Figure 1a**). The small model enables extensive ISA for genome-wide interpretation of the motif syntax underlying TF cooperativity, while maintaining state-of-the-art prediction accuracy for the effect size of genetic variants. We identify pairwise additivity (independence) as the baseline of the majority of TF pairs, and formally define a TF cooperativity score to systematically quantify deviations from the baseline model. We demonstrate that within the deep learning framework, redundancy and synergy lie on opposite sides of the same nonlinearity spectrum. Although additivity is prevalent, cooperativity is critical to explain the emergent properties of promoters and enhancers. TFs that primarily act in a redundant manner are more ubiquitously expressed, more important for promoter activity, and more conserved in promoters compared to enhancers. Conversely, TFs that are primarily synergistic with partner TFs are more likely to physically interact, require closer motif distance, are more cell-type specific, and more important for enhancer activity. Our analytical framework has major implications for future modeling and interpretation of nonlinear gene regulatory effects.

**Figure 1.**
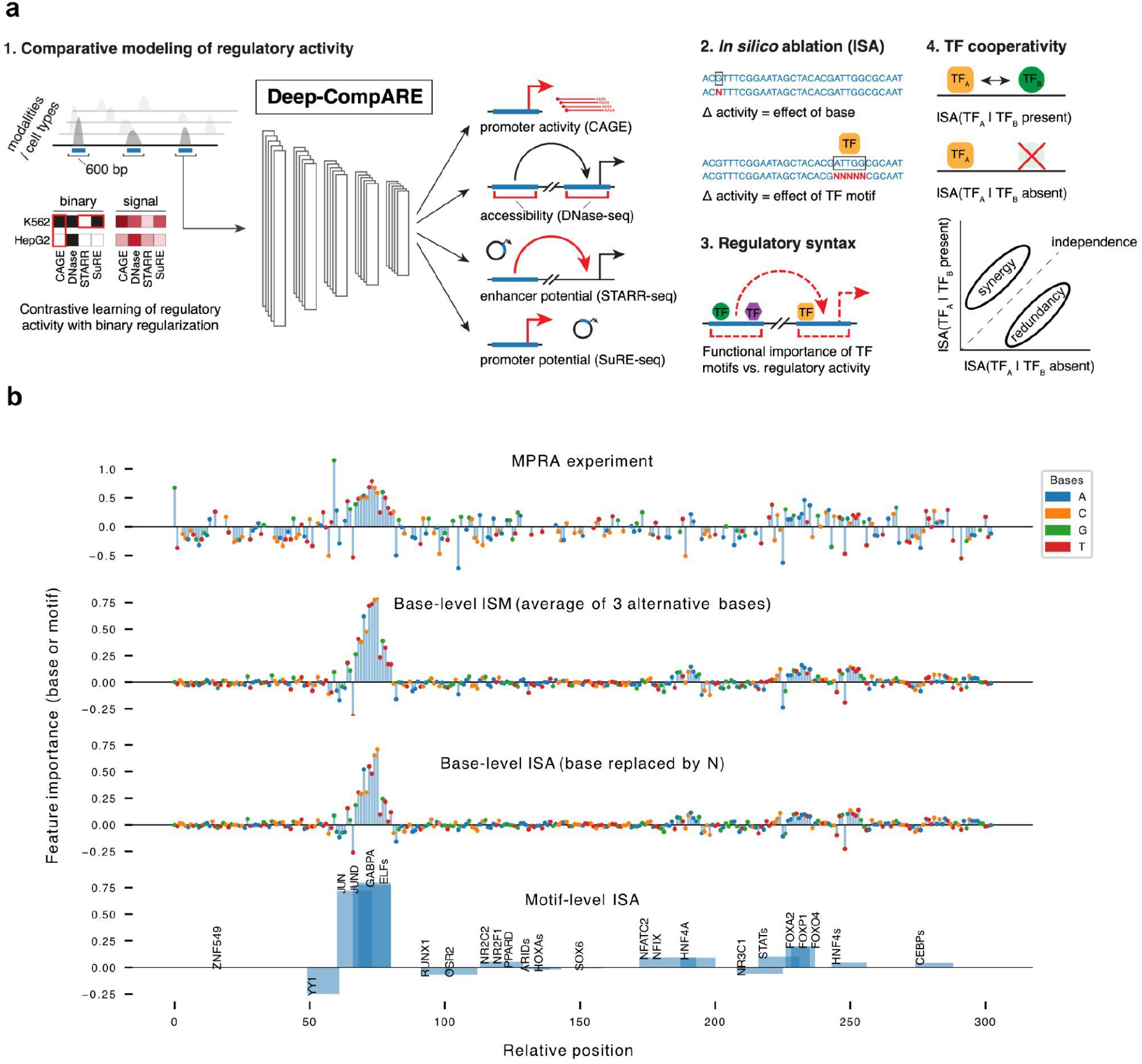
Deep comparative modeling of regulatory activity from DNA sequence reveals TF motif syntax. **a:** Schematic overview of DeepCompARE (1) and the computational framework based on *in silico* ablation (ISA, 2) to study TF motif syntax (3) and TF cooperativity (4). **b:** From top to bottom: MPRA mutagenesis data^32^ reflecting the effect size (vertical axis) of mutating the base of each position (horizontal axis) in the F9 promoter (first track), ISM-derived importance scores from DeepCompARE SuRE-seq K562 track (vertical axis) from averaging of all 3 alternative bases for each position (second track), corresponding ISA-derived importance scores (vertical axis) from masking the base in each position (third track), and motif-level ISA importance scores (vertical axis) derived from masking all putative TF binding sites^36^ in the promoter sequence, one by one (fourth track).

## Results

### Deep comparative modeling of regulatory activity from DNA sequence with DeepCompARE

To model the grammar of gene regulatory elements, we trained a BPNet^22^-inspired lightweight deep convolutional neural network model for comparative analysis of regulatory elements (DeepCompARE). DeepCompARE was trained to predict from DNA sequence the average log-transformed signal intensity of CAGE (promoter activity, this study), DNase-seq^29^ (chromatin accessibility), STARR-seq^29^ (enhancer potential), and SuRE-seq^30^ (promoter potential) data within regulatory elements in K562 erythroleukemia and HepG2 hepatoblastoma cells. For regularization purposes, DeepCompARE was trained to also predict a binarized label (classification) for each data modality. The size of the resulting model, with fewer than 440k parameters, a 1/500th of that of the state-of-the-art model Enformer^31^, enables extensive *in silico* perturbations and TF binding site ablation to explore regulatory syntax and TF cooperativity (**Figure 1a**). As a result of data-dependent regularization (**Supplementary Figure 1a-c**), careful data labeling and filtering (**Supplementary Figure 1d**), contrastive learning (**Supplementary Figure 1e**) and intentional reduction of parameters (Methods), the regression branches of DeepCompARE achieved a high correlation between predicted and true signal intensity on held-out test data (mean Pearson *R* = 0.76, 0.67-0.84 across tracks; **Supplementary Figure 1f**).

To further evaluate the predictive performance of DeepCompARE, we benchmarked zero-shot predictions of the effect sizes of single nucleotide variants (SNVs) on regulatory activities against Enformer. DeepCompARE performed better or on par with Enformer on several disease-relevant promoters and enhancers according to MPRA saturation mutagenesis data^32^ (**Supplementary Figure 2a**). Closer inspection of the *F9* gene promoter revealed a high similarity (Pearson *R* = 0.67) between base-level MPRA effect sizes and base-level importance scores derived from *in silico* mutagenesis^33^ (ISM) of the DeepCompARE SuRE-seq track for HepG2 (**Figure 1b**).

Albeit accurate, ISM requires intensive computation, with the number of predictions scaling exponentially with the length of the investigated sequence, severely impeding investigations of motif syntax. To reduce computation and avoid unintended creation of novel motifs during ISM (**Supplementary Figure 2b**), we developed *in silico* ablation (ISA), revealing the importance of a base or a motif by masking the corresponding sequence information. Hence, ISA scores reflect perturbation effect sizes on predicted activity. Importantly, base-level ISA closely resembled base-level ISM (Pearson *R* > 0.94; **Figure 1b**; **Supplementary Figure 2c**), and motif-level ISA scores were similar to gradient-derived motif importance^34^ (Pearson *R* > 0.75; **Supplementary Figure 2d**) and highly similar to the averages of ISM scores from complete combinatorial perturbations of motifs (Pearson *R* = 0.94±0.02; **Supplementary Figure 2e**).

Consequently, motif-level ISA scores accurately reflected the experimentally derived base-level importances of the *F9* gene promoter and revealed high importance of an ETS binding site located ~250bp upstream of the *F9* gene transcription start site (TSS) (**Figure 1b**). Furthermore, DeepCompARE base-level ISA scores correctly interpreted the gain-of-function mutation chr1:109274968 G>T, which creates a *de novo* CCAAT-Enhancer-Binding Protein (C/EBP) binding site and increases *SORT1* expression in the liver^35^. DeepCompARE systematically elevated the base-level ISA score of most bases within the novel C/EBP binding site (**Supplementary Figure 2b**).

### In silico ablation reveals the importance of cell- and modality-specific TFs

To assess the importance of each TF on regulatory activity, we calculated motif-level ISA scores for all putative binding sites^36^ of expressed TFs (TFBSs), based on curated motifs, across a strict set of 12,614 promoters and 15,238 putative enhancers for K562 (10,450 and 16,775 for HepG2, respectively). Overall, TFBS ISA scores were highly correlated across modalities (Pearson *R* from 0.60 to 0.95; **Supplementary Figure 3a**). Furthermore, TFBS ISA scores collected in promoters showed higher consistency across cell types than those of enhancers, indicating that the latter are more responsible for cell-type specific transcriptional regulation (Pearson *R* = 0.91 for promoters vs 0.80 for enhancers; **Supplementary Figure 3b**). Despite a general ISA concordance across tracks, DeepCompARE also highlighted track-specific factors. For instance, key regulators of liver function like C/EBP and Hepatic Nuclear Factor (HNF) family TFs were found to be specifically important for HepG2 tracks, whereas well-known regulators of hematopoiesis and leukemogenesis, including GATA family TFs and members of the Signal Transducer and Activator of Transcription (STAT) TF family, were highlighted by K562 tracks (**Figure 2a**). Other TFs displayed specific association with regulatory activity. Among these are Forkhead box O (FOXO) family TFs, specifically enriched among TFs important for enhancer activity in K562, and Nuclear Transcription Factor Y family (NFY) TFs and Krüppel-Like Factor (KLF) TFs, specifically important for promoter activity in K562 (**Figure 2b**).

**Figure 2.**
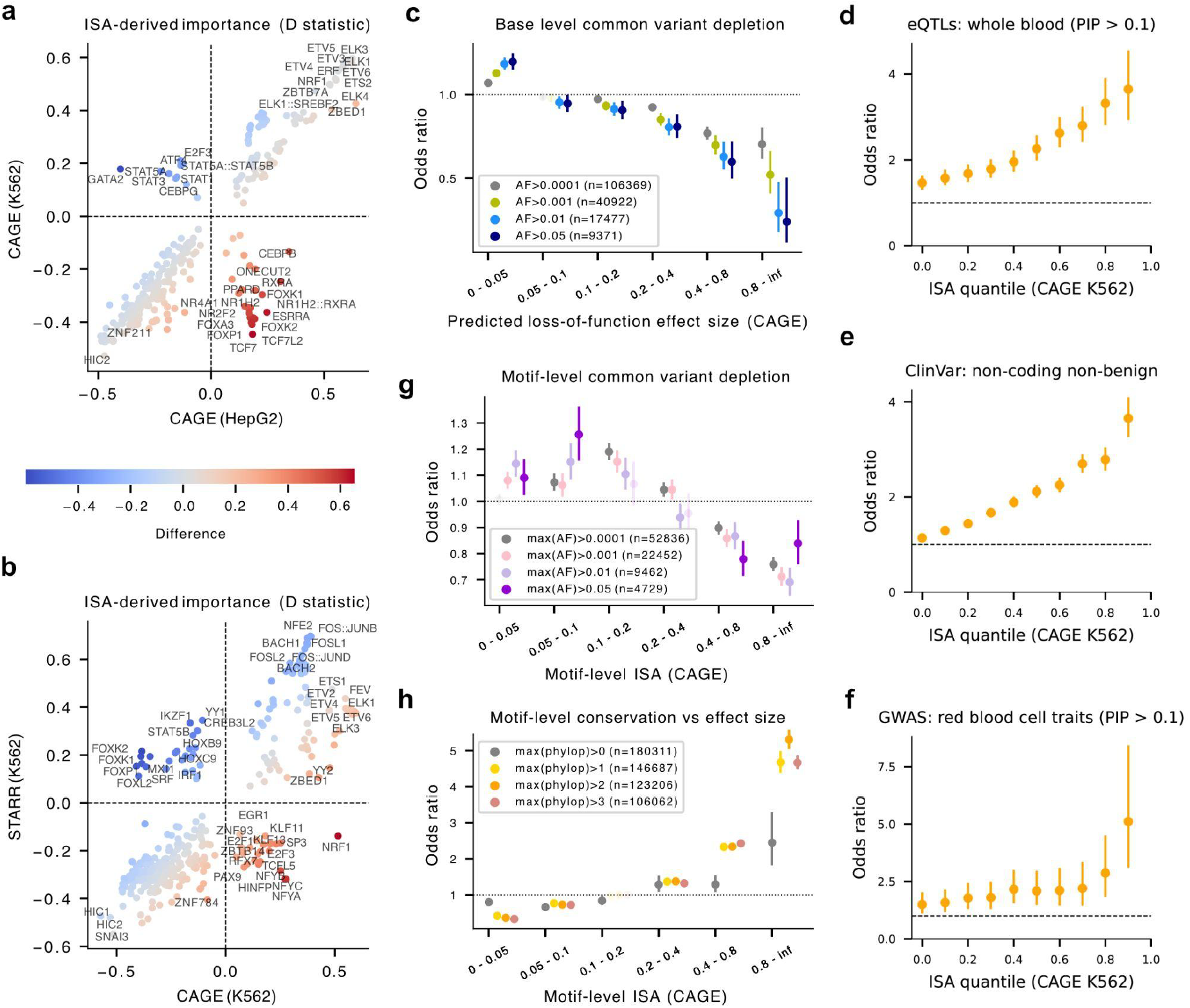
Functional relevance of DeepCompARE-derived *in silico* ablation scores. **a:** Differences in ISA-derived TF motif importance from DeepCompARE CAGE tracks for K562 (vertical axis) and HepG2 (horizontal axis), as measured by signed Kolmogorov-Smirnov test D statistics (Methods). Positive values indicate a higher relative importance compared to other TFs. **b:** As **a**, but comparing the inferred importance of TFs for CAGE signal (horizontal axis) versus STARR-seq signal (vertical axis) in K562 cells. **c:** Predicted loss-of-function effect size on K562 CAGE track (horizontal axis), versus enrichment (odds ratio, vertical axis) of variants from gnomAD^38^, thresholded by minor allele frequency (MAF). Transparent dots represent P >= 0.05. Solid dots represent P < 0.05. Error bar: 95% CI. **d-f**: Enrichment of (**d**) fine-mapped whole-blood eQTLs^39^ (PIP > 0.1), (**e**) ClinVar non-coding and non-benign variants^40^, and (**f**) fine-mapped GWAS variants for red blood cell traits^41^ (PIP > 0.1), versus common variants. **g-h:** As **c**, but considering motif-level ISA scores (horizontal axis) versus (**g)** the enrichment of variants (vertical axis) thresholded by the maximum MAF across the motif and (**h**) the enrichment of the maximum phyloP score^42^ across the motif (vertical axis).

Regardless of cell type, ETS family TFs were predicted to be the most important for promoter activity, whereas Activator Protein 1 (AP-1) factors, including JUN, FOS and ATFs, were predicted to be most important for enhancer activity (**Figure 2b**). CTCF, a key regulator of chromatin architecture, was specifically highlighted by DNase tracks and found to be of low importance for predicting CAGE signal (**Supplementary Figure 3c**), in line with reduced transcription initiation at CTCF-bound boundary elements^37^.

### In silico ablation scores reflect mutational constraints and evolutionary conservation

We next assessed the utility of DeepCompARE ISA scores to gain functional insights into genomic sequences. We observed that common variants^38^ were significantly depleted (MAF > 1%, odds ratio (OR) = 0.63, P = 1.43e-13 for 0.4 ≤ ISA < 0.8; Fisher exact test, **Figure 2c**, bright blue) among SNVs predicted to have a loss-of-function effect, and the depletion increased by effect size (OR = 0.29, P = 2.3e-9 for base-level ISA ≥ 0.8, Fisher exact test). A reciprocal enrichment was observed for variants with low predicted effect size. This association was largely reproducible across cell types, regulatory elements and modalities, with CAGE and SuRE-seq tracks best capturing the mutational constraints at promoters, and STARR-seq and SuRE-seq tracks best reflecting those of enhancers (**Supplementary Figure 4**). This indicates that variants with large regulatory effect sizes are less tolerated in the human population and that DeepCompARE base-level ISA scores capture potentially deleterious variants. In support, variants with larger effect sizes were more likely fine-mapped eQTLs^39^, possibly pathogenic variants^40^, or fine-mapped GWAS variants^41^ (**Figure 2d-f; Supplementary Figure 3d-f**).

Among TF activators (motif-level ISA > 0), we similarly observed that TFBSs with large effect sizes were less likely to harbor common variants^38^ (e.g., OR 0.93, P = 0.02 for 0.2 ≤ ISA < 0.4 and OR 0.69, P = 1.7e-22 for ISA > 0.8, Fisher exact test; **Figure 2g**) and more likely to be conserved^42^ (e.g., considering motifs with maximum phyloP > 3: OR 1.32, P = 1.03e-106 for 0.2 ≤ ISA < 0.4; OR 4.67, P < 2.2e-308 for ISA > 0.8, Fisher exact test; **Figure 2h**). These observations are reproducible across cell types, regulatory elements and modalities (**Supplementary Figure 5**), emphasize the predictive value of DeepCompARE-derived ISA scores, and suggest that these can be used to gain insights into the functional roles of TFBSs.

### Pairwise independence between distant TF binding sites

Having derived a method to efficiently assess the importance of individual TFBSs for regulatory activity, we next sought to understand the interaction effects between pairs of TFBSs as a proxy for TF cooperativity. We focused specifically on TFBSs with an inferred activating effect (motif-level ISA > 0), and used a top-down approach to model interaction effects as the difference in ISA of a TFBS with or without a partner TFBS, with the rest of the sequence left intact (**Figure 3a**). This strategy allows capturing pairs of TFBSs that act in an independent or in a cooperative manner. If two TFBSs independently contribute to regulatory activity, their ISA scores should not be affected by the presence or absence of each other, thus their effects are additive (Methods, **Figure 3a**). TFBS interaction can therefore be observed through deviation from perfect additivity. We note that, since DeepCompARE predicts log-transformed signals, an additive effect on the model’s predictions corresponds to a multiplicative effect on raw signals.

**Figure 3.**
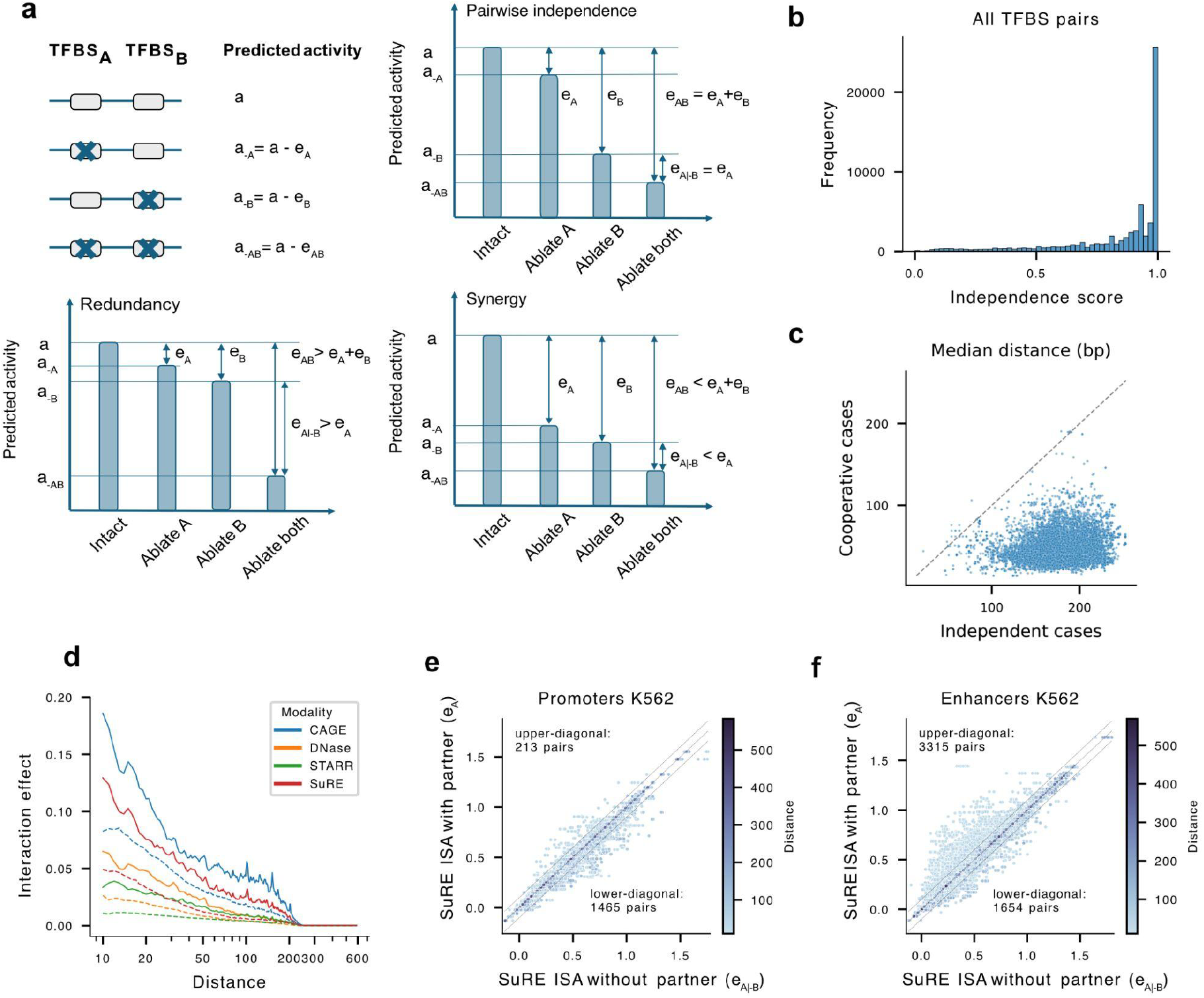
Two modes of TF cooperativity, redundancy and synergy, explain deviation from pairwise TF independence. **a:** Schematic of the top-down approach used to determine the cooperativity between pairs of TFBSs (here denoted TFBS_A_ and TFBS_B_) based on motif-level ISA. Notation: the activity of the intact sequence (a), the activity with TFBS_A_ ablated (a_-A_), and the activity with both TFBS_A_ and TFBS_B_ ablated (a_-AB_). The ISA score (effect) of TFBS_A_ with TFBS_B_ present (e_A_) is the difference between a and a_-A_. The effect of TFBS_A_ in the absence of TFBS_B_ (e_A|-B_) is the difference in activity between sequence with TFBS_B_ ablated and sequence with both TFBS_B_ and TFBS_A_ ablated. Two TFBSs are pairwise independent (top right) if the effect of removing TFBS_A_ is unaffected by the presence or absence of TFBS_B_, resulting in additivity. Redundancy (bottom left) occurs when ablating TFBS_A_ causes a larger drop in activity when TFBS_B_ is absent. Synergy (bottom right) occurs when ablating TFBS_A_ causes a larger drop in activity when TFBS_B_ is present. **b:** Histogram of independence scores for all TFBS pairs in K562. **c:** Median distance between independent (horizontal axis) and cooperative (vertical axis) TFBS pair instances for each TF pair in K562. **d:** TF partner effect (|e_A_ - e_A|-B_|, vertical axis) as a function of TFBS distance. **e:** SuRE-seq-derived motif-level ISA scores of all FOS::JUN binding sites on K562 promoters, comparing effects without the partner (e_A|-B_, horizontal axis) versus with the partner (e_A_, vertical axis). Off-diagonal cases (upper and lower) are defined by an ISA difference of at least 0.1 (|e_A_ - e_A|-B_| ≥ 0.1). Dashed lines represent the 0.1 threshold. **f:** As **e**, but for FOS::JUN binding sites on K562 enhancers.

We defined independent cases as those with negligible interaction effects (absolute difference < 0.01; Methods) and cooperative cases (dependent) as those with large interaction effects (absolute difference > 0.1). We calculated an independence score defined as the fraction of independent cases among all co-occurring TFBS pairs for each individual TF pair. Around half of TF pairs (53.94% for HepG2 and 58.83% for K562) behaved in a pairwise independent manner in more than 90% of instances (**Figure 3b; Supplementary Figure 6a**). Interestingly, for >99% of TF pairs with both independent and cooperative TFBS pair instances, the median genomic distance between TFBSs in cooperative pairs was shorter than between those of independent pairs (**Figure 3c; Supplementary Figure 6b**). Regardless of modality, the average influence of a TFBS partner decreases as distance increases (**Figure 3d**). We note that, beyond 255 bp, the influence of partner TFBSs is zero, which reflects the receptive field of DeepCompARE (Methods). However, a model with a larger receptive field of 511 bp showed a similar decline (**Supplementary Figure 6c**). We observed similar yet milder trends by replacing the partner TFBSs with random sequences of similar length (**Figure 3d**, dashed lines). Together, these results indicate independent, additive behavior for the majority of TFBS pairs.

### Redundancy and synergy explain opposite modes of TF cooperativity

To understand the cooperativity between TFBS pairs in close proximity, we next assessed, for each TF or TF composite, the effect sizes of all its binding sites on promoters and enhancers, in the presence versus absence of another TFBS (agnostic to partner TF identity). This revealed an association between TFBS cooperativity and regulatory function. For example, combinatorial motif-level ISA of FOS::JUN TFBSs paired with any other TFBS revealed deviations from pairwise independence in opposite directions at promoters and enhancers: FOS::JUN binding sites tend to become more important upon partner TFBS loss at promoters (1,465 lower diagonal cases vs. 213 upper diagonal cases, **Figure 3e)**, whereas at enhancers, the majority of FOS::JUN binding sites become less functional in the absence of partner TFBSs (1654 lower diagonal cases vs. 3315 upper diagonal cases, **Figure 3e,f**).

To explain the TFBS cooperativity (deviation from pairwise independence) indicated by upper and lower diagonal biases in **Figure 3e,f** (bottom), we propose two modes of cooperativity: redundancy and synergy. Redundancy occurs when two TFBSs have functional overlap: a TFBS becomes less important (causes smaller drop in activity upon ablation) when its partner is present. Synergy, on the other hand, occurs when the absence of a partner TFBS impedes the TFBS from fully exerting its function: a TFBS becomes less important when its partner is lost (**Figure 3a**; **Supplementary Figure 6d**).

To quantify the cooperativity between each pair of TFs, as indicated by the effects of their binding sites, we calculated the differences between TFBS effects with and without partner TFBSs across all instances of TFBS pairs in promoters or enhancers. Next, we defined a synergy score of a TF pair as the fraction of synergistic cases among all cooperative cases, weighted by interaction strength, and similarly, a redundancy score as the weighted fraction of redundant cases (Methods). By definition, a synergy score of 0 for a TF pair means that all their TFBS pair instances in regulatory elements display redundant cooperativity (i.e., redundancy score = 1).

As expected (**Figure 3c**), cooperative TF pairs had, in general, more TFBSs closer to one another than those of independent pairs (**Figure 4a**; **Supplementary Figure 7a**). Furthermore, TFBSs of synergistic TF pairs (synergy score > 0.7 or redundancy score < 0.3) were less distant than TFBSs of redundant pairs (synergy score < 0.3 or redundancy score > 0.7) (**Supplementary Figure 6e**,**f**), indicating that synergistic TFs may achieve cooperativity through a mechanism that requires proximity, e.g., via a direct protein-protein interaction (PPI). As posited, we observed that TF pairs supported by PPI data^43,44^ had significantly higher synergy scores (P < 2.4e-193, two-sided Mann-Whitney U test) and lower independence scores (P < 2.2e-308, two-sided Mann-Whitney U test) than those lacking PPI support (**Figure 4b,c**). Furthermore, TF pairs for which both partners interacted with a common third factor tended to have higher synergy scores than those that lacked a common partner (P = 1.1e-9, two-sided Mann-Whitney U test; **Figure 4d**), indicating that synergy can also be mediated by a third factor that interacts with both TFs. This trend is particularly apparent for co-interactions with Mediator (**Figure 4d**), the SWI/SNF chromatin remodelling complex, RNA polymerase II, and most general TFs (GTFs) (**Supplementary Figure 7**).

**Figure 4.**
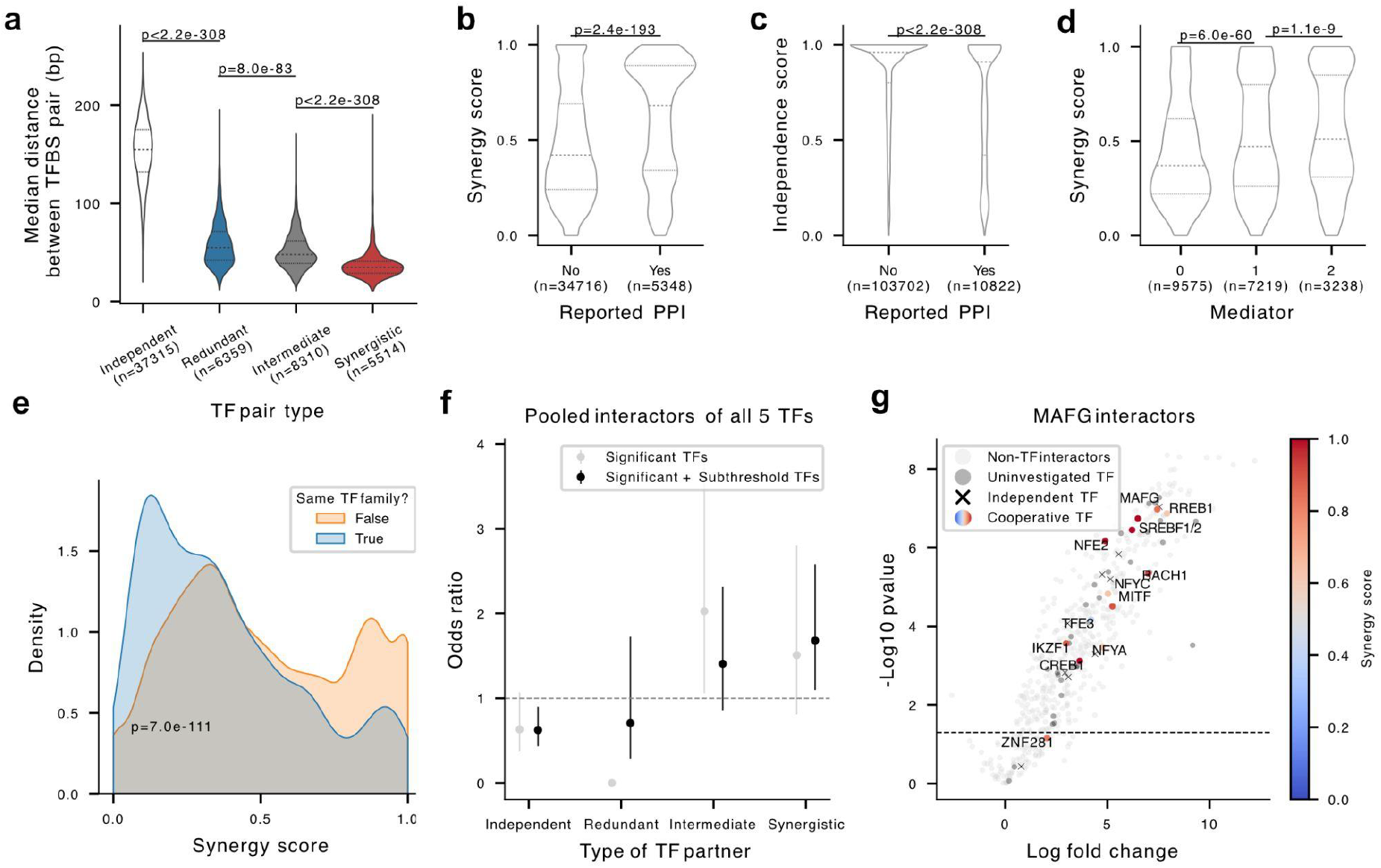
Synergistic TF pairs indicate protein-protein interaction. **a:** Distribution of median genomic distance between TFBS pairs for independent, redundant, intermediate, and synergistic TF pairs. **b:** Distribution of TF pair synergy score, grouped by whether the TF pair has PPI record in public databases (STRING^43^ or InWeb^44^). **c:** Distribution of TF pair independence score, grouped by whether the TF pair has PPI record in public databases. **d:** Distribution of TF pair synergy score, grouped by the number of mediator-interactors in the TF pair. **e:** Distribution of TF pair synergy score, grouped by whether the TF pair belong to the same TF family. **f:** Enrichment (odds ratio) of independent, redundant, intermediate, and synergistic TF partners captured by IP-MS interaction experiments. Black dots are the odds ratios calculated using all TF interactors including the TFs that failed stringent IP-MS filter (Methods), whereas gray dots consider only the TFs that pass the filter. Error bar: 95% CI. **g:** Volcano plot for MAFG interactors, with log-fold change of TF bait over IgG displayed on the horizontal axis and −log_10_ p-value from a two-tailed two-sample moderated t-test displayed on the vertical axis (Methods). Light gray dots: Protein interactors that are not TFs. Dark gray dots: TFs uninvestigated by combinatorial ISA due to lack of high-quality predicted binding site locations in JASPAR database. Black crosses: TF partners predicted to act independently from the TF bait. Color of dots indicate the synergy score of cooperative TFs.

To investigate a possible mechanistic relationship between synergistic TFs through PPI, we selected 5 TFs (MAFG, BACH1, IKZF1, RREB1, RFX5) with high synergy scores and few reported PPIs in public databases^43,44^ and performed immunoprecipitation-mass spectrometry (IP-MS) to capture their TF interactors in K562 cells. Reassuringly, our experiments revealed novel TF interactors correctly indicated by high synergy score (**Figure 4g**; **Supplementary Figure 8**). Overall, TF partners captured by IP-MS tended to have higher synergy scores and lower independence scores than non-interactors, and the likelihood of TF interaction increased from independent to cooperative TFs, and among cooperative TFs from redundant to synergistic TFs, with intermediate TFs (0.3 < synergy score < 0.7) falling in between (**Figure 4f**).

While synergistic TFs displayed a higher probability of being mediated through PPI (**Figure 3b-d,f-g**), TF pairs of the same TF family tended to have higher redundancy scores (lower synergy scores) than TF pairs from different families (P = 1.3e-25, two-sided Mann-Whitney U test; **Figure 4e**). Hence, compensation through functional similarities among TFs from the same family binding to proximal motifs provides a mechanistic interpretation of redundant TFs.

### Functions of redundant and synergistic TFs

We next explored similarities between TFs as measured by their synergy scores with partner TFs. Interestingly, TFs formed apparent groups that were consistently either redundant or synergistic with the majority of their partners (**Figure 5a**; **Supplementary Table 2**,**3**). For example, the KLF family TFs displayed redundancy towards the majority of TFs. In contrast, AP-1-related TFs were largely synergistic with partners.

**Figure 5.**
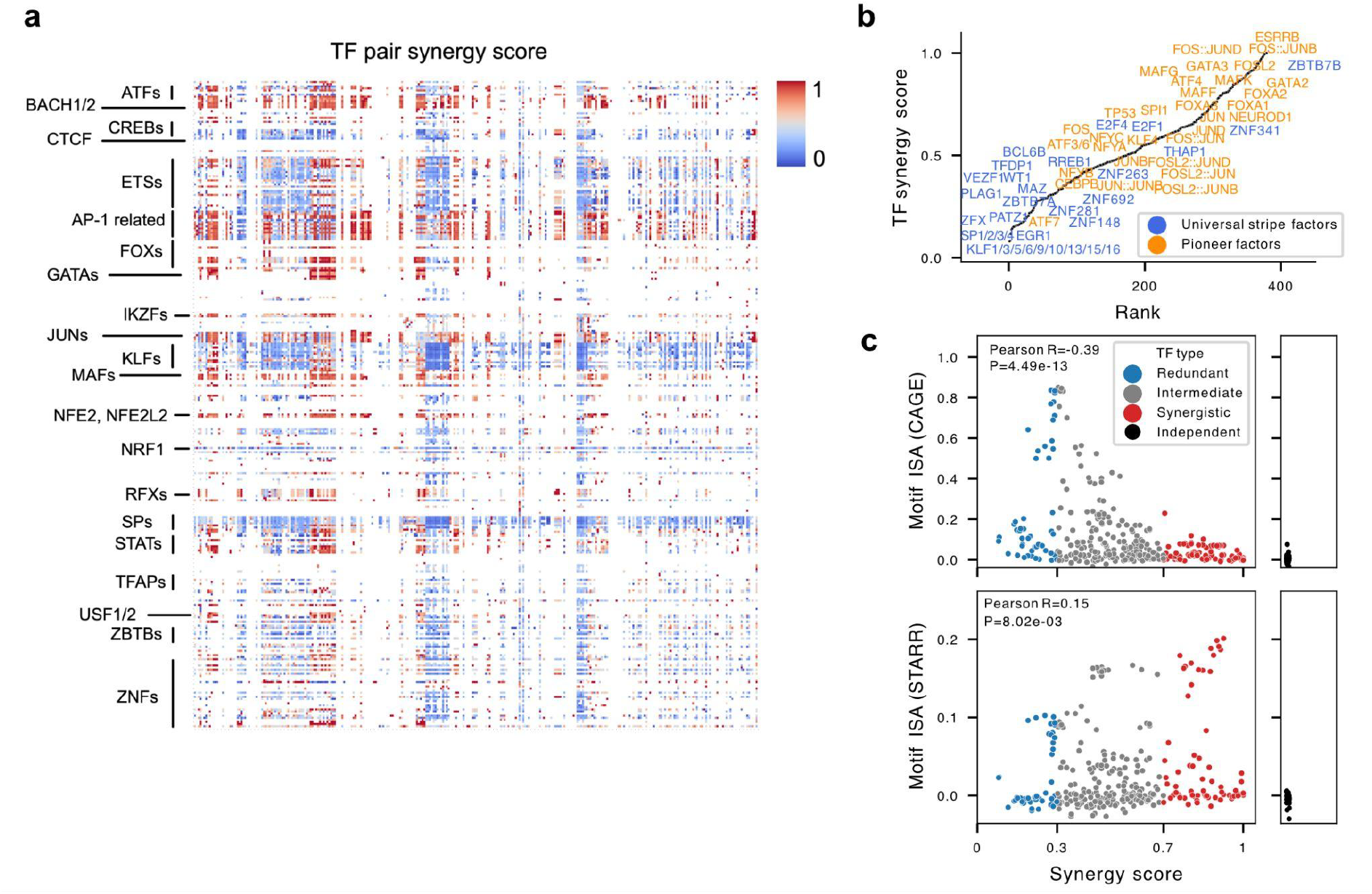
Properties of TF-level synergy score. **a:** Heatmap (symmetric) of TF pair synergy scores, sorted by TF name. **b:** Ranked TF synergy score with annotations of universal stripe factors (blue) and pioneer factors (orange) for K562. **c:** Synergy score (horizontal axis) is negatively correlated with CAGE-derived ISA score (vertical axis, top panel), and positively correlated with STARR-seq-derived ISA score (vertical axis, bottom panel). ISA scores of independent TFs are shown on the right panel due to lack of synergy score.

To reveal the general cooperative behavior of TFs, we categorized each TF according to a TF level synergy score regardless of its partners (Methods; redundant TFs: TF-level synergy score < 0.3, synergistic TFs: TF-level synergy score > 0.7; **Supplementary Figure 9a**). This approach identified 58 redundant TFs and 96 synergistic TFs for K562 (53 and 99 for HepG2, respectively; **Supplementary Table 4**,**5**). Redundant TFs include the ETS, KLF, and Specificity Protein (SP) family TFs: broadly expressed TFs that bind redundantly to regulatory elements^20,45,46^. Synergistic TFs include GATA, FOX, STAT, and AP-1 TFs, which contribute to highly specialized functions including cell fate determination and stress response, and are in general more cell-type specific (HepG2: Pearson R = 0.31, P = 1.6e-8; K562: R = 0.17, P = 1.2e-3; **Supplementary Figure 9b**). TFs with Forkhead, Homeo, and nuclear receptor DNA binding domains were found to be most synergistic, whereas TFs with C2H2 zinc finger, Rel and Ets domains were most redundant (**Supplementary Figure 9c**). In support, we noted that Universal Stripe Factors, which are ubiquitously expressed and bind redundantly to increase chromatin accessibility^21^, indeed had a significantly lower synergy score (i.e., higher redundancy score) than other TFs (P = 1.6e-11, Mann-Whitney U test). In contrast, pioneer factors^47^, which are critical for cell type specific transcriptional regulation^7^, displayed higher synergy scores than non-pioneer factors (P = 0.04, two-sided Mann-Whitney U test) (**Figure 5b**; **Supplementary Figure 9d**).

Lastly, we explored the relationship between the TF-level synergy score and regulatory function. Interestingly, synergy scores are negatively correlated with CAGE-derived motif-level ISA scores (Pearson R = −0.49 for HepG2 and R = −0.39 for K562), but positively correlated with STARR-seq-derived scores (Pearson R = 0.18 for HepG2 and R = 0.15 for K562; **Figure 5c**; **Supplementary Figure 9e**). Hence, redundant TFs have a larger influence on promoter activity, whereas synergistic TFs drive enhancer activity. This is consistent with the generally high GC content of redundant TFBSs and promoters, and the lower GC content associated with enhancers and synergistic TFBSs (**Supplementary Figure 9f)**. In summary, synergistic TFs predominantly influence enhancer activity, while redundant TFs exert stronger effects at promoters.

### Synergy and redundancy have different contextual importance

To characterize genome-wide rules of TF cooperativity, we next focused on DNase hypersensitive sites (DHSs)^48^, comprising 38,492 promoter-proximal and 193,806 promoter-distal DHSs for HepG2 (40,785 and 269,572 DHSs for K562, respectively). A synergy score calculated from the DHS set largely agreed with that derived from the smaller set of enhancers and promoters (For HepG2, Pearson R = 0.83 at TF pair level and 0.82 at TF level; For K562, R = 0.81 at TF pair level and R = 0.77 at TF level; **Figure 6a**; **Supplementary 10a-c**), demonstrating that our analytical framework to study cooperativity is robust. To correct for systematic inflation caused by the greater presence of promoter-distal regions (putative enhancers) in the DHS set, we calibrated the synergy score. This calibration ensured that 68% of redundant TFs and 70% of synergistic TFs could be accurately retrieved through DHS analysis (for K562: 67% or redundant TFs and 60% of synergistic TFs were retrieved).

**Figure 6.**
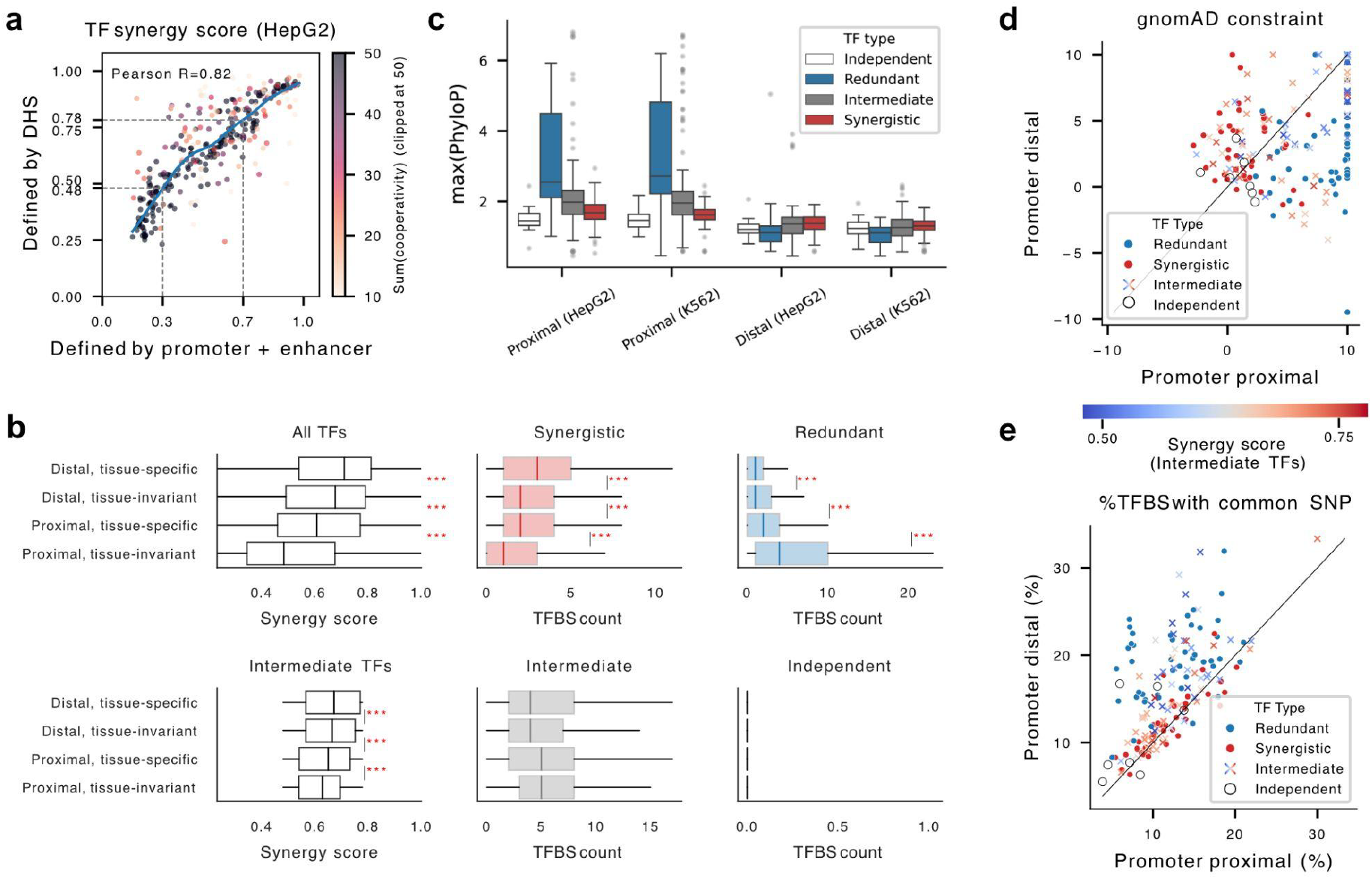
Differential contextual importance of redundant, intermediate and synergistic TFs. **a:** Correlation of TF-level synergy score defined by promoter and enhancer (horizontal axis), versus synergy score defined by DHSs (vertical axis). **b:** Left panel: Distribution of TF-level synergy score of all TFs (top) and intermediate TFs (bottom), grouped by region type. The middle and right panels show the distributions of the number of binding sites per regulatory element for synergistic (middle) and redundant (right) TFs, grouped by region type. Box plot: the box represents the interquartile range (IQR), with the median shown as a line inside the box. Whiskers extend to 1.5 times the IQR, and outliers beyond this range are not shown. ***: P≤1e-3, **: 1e-3≤P<1e-2, *:1e-2≤P<0.05. **c:** Distribution of maximum phyloP score over motifs for each type of TF, within each group of regulatory elements. **d:** gnomAD constraints for binding sites of each TF (see Methods), calculated from promoter-proximal (horizontal axis) or promoter-distal (vertical axis) regions. Each dot represents one TF. **e:** As **d**, but comparing the fraction of TFBSs harboring at least one common SNP in promoter-proximal (horizontal axis) versus promoter-distal (vertical axis) regions. Redundant and synergistic TFs are shown as blue and red dots, respectively. Intermediate TFs are colored by their synergy score.

A focus on the larger DHS set allowed a more systematic comparison of TF cooperativity across different regulatory elements with respect to genomic location and expression specificity. We observed a gradual decrease of TF-level synergy scores from tissue-specific to tissue-invariant distal regions and from tissue-specific to tissue-invariant proximal regions. This was supported by a similar decline in the number of TFBSs for synergistic TFs at proximal and tissue-invariant regions and a gradual increase in the number of TFBSs for redundant TFs. The TFBS count for independent or intermediate TFs showed no similar trends, yet the median synergy score of intermediate TFs displayed a decreasing trend, indicating that the association between cooperativity and cell-type-specific control extends beyond the strictly defined redundant and synergistic TF categories (**Figure 6b**; **Supplementary Figure 10d**). Interestingly, we observed a similar gradient in synergy score and number of TFBSs for synergistic TFs along strong enhancers (E1), weak enhancers (E2), potentially enhancer-containing strong promoters (P2), and weak promoters (P1) defined by MPRA data^49^ (**Supplementary Figure 10e**), consistent with our observation that synergistic TFs are the major regulators of enhancer activity.

To further investigate the contextual functional importance of synergistic and redundant TFs, we assessed their TFBS conservation (as measured by the max Zoonomia phyloP score^42^) across DHSs. We noted that the relative conservation levels of redundant and synergistic TFs were reversed in different groups of regulatory elements (**Figure 6c**). In promoter-proximal regions, the binding sites of redundant TFs are more conserved than those of synergistic TFs (P = 3.6e-14 for HepG2, P = 4.0e-12 for K562, two-sided Mann-Whitney U test), while the opposite trend is true in distal regions (P = 1.1e-7 for HepG2 and P = 6.1e-6 for K562, two-sided Mann-Whitney U test). Hence, while promoter-distal TFBSs are overall less conserved compared to promoter-proximal ones, those of synergistic TFs showed a relatively higher conservation, emphasizing their higher importance for enhancer activity. Consistent with the elevated importance of synergistic TFs in distal regions, we observed a negative correlation between synergy score and maximum TFBS phyloP score in promoter-proximal regions (R = −0.49, P = 1.3e-20 for HepG2; R = −0.43, P = 2.1e-16 for K562), while the correlation was positive in distal regions (Pearson R = 0.15, P = 6.1e-3 for HepG2; R = 0.18, P = 8.7e-4 for K562). A notable exception to these observations is CTCF, which displayed high conservation in both distal and proximal regions, reflecting its architectural role in promoter-distal regions^50^ (**Supplementary Figure 10f**).

In support of the opposite contextual importance of synergistic and redundant TFs, analysis of human genomic constraint (gnomAD gnocchi z score^38^ repurposed for TFBSs, Methods), revealed that 65% of synergistic TFs had more constrained binding in promoter-distal than in promoter-proximal regions, whereas 92% of redundant TFs showed the reversed pattern (91% and 88% for K562, respectively) (**Figure 6d**; **Supplementary Figure 10g**). In addition, 92% of redundant TF binding sites harbor common variants (MAF > 0.1%) more frequently in promoter-distal than promoter-proximal regions, whereas only 41% synergistic TFs show such bias (88% and 33% for K562, respectively). Both trends extend to intermediate TFs (**Figure 6e**; **Supplementary Figure 10h**). In conclusion, synergy and redundancy exhibit context-dependent functional importance at enhancers and promoters, respectively.

## Discussion

We present DeepCompARE, an accurate, lightweight, multitasking convolutional neural network, along with *in silico* ablation (ISA), a highly computationally efficient interpretation method. ISA exponentially reduces the computational cost of calculating motif importance compared to ISM, allowing us to systematically quantify TF motif importance as well as the pairwise cooperativity between TF motifs genome-wide.

We found that while around half of the investigated TF pairs behave additively (independently), the remaining pairs deviate systematically from the additive baseline. These deviations can be explained by functional redundancy and synergy, which we rigorously define using a top-down approach. We note that previous studies have described similar phenomena using different terminology^25,26,28^. The most common alternative has been to take a bottom-up viewpoint. Since the addition of a functionally redundant TFBS will not further increase the overall activity, the effect of the TFBS pair as a whole will be smaller than the sum of their individual effects, a phenomenon often termed ‘sub-additivity’, or ‘saturation’. Likewise, TF synergy is often termed ‘super-additivity’. However, since the comparison between effect-of-sum and sum-of-effect reverses the direction between top-down and bottom-up viewpoints (**Supplementary Figure 6d**; Methods), sub- and super-additivity can become incomparable across studies.

We found that TF cooperativity helps explain fundamental differences between promoters and enhancers^4^. Promoters are generally less responsive to environmental stimuli, a property essential for maintaining homeostasis, whereas enhancers are more responsive, enabling adaptation and regulatory specificity (**Supplementary Figure 3b**). At the low end of the responsiveness spectrum, housekeeping promoters are robustly active, even in the presence of DNA sequence mutations and cell-type-specific variation in TF protein abundance. Such robustness arises from redundancy: as long as a sufficient subset of TFBSs remain capable of recruiting TFs with necessary activating functions, promoter activity will remain stable. TF redundancy thus resolves a paradox: although housekeeping gene bodies are more conserved than those of tissue-specific genes, their promoters are less conserved^37,51^. Our analysis reveals that promoter-proximal regions with tissue-invariant activity exhibit higher redundancy than tissue-specific ones, explaining the higher mutational tolerance of housekeeping promoters (**Figure 6c**).

At the high end of the responsiveness spectrum, promoter-distal regions with tissue-specific activity exhibit the strongest TF synergy. We suspect that such synergy plays a key role in the temporal and spatial specificity of enhancer activation. Because multiple TFs encode and integrate diverse internal and external cues, the absence of even a single TF may indicate that a required condition is unmet. As a result, the enhancer remains partially inactive—a safeguard enforced by synergistic regulation. In support, enhancers critical for organ development are enriched for low-affinity degenerate TFBSs to enforce TF co-binding, whereas mutations that increase binding affinity of individual TFs will allow TFs to bind and work alone, reduce synergistic operation, diminish enhancer specificity, and cause inappropriate activation in ectopic tissues^52^. We further speculate that the synergy level of a regulatory element can be a proxy for its syntactical rigor. For example, promoter-distal regions are more syntactically rigorous than promoter-proximal ones, whereas cell-type-specific regulatory elements are more syntactically rigorous than ubiquitous ones.

Apart from improving the robustness of gene regulation, we speculate that TF cooperativity can enable flexible fine tuning of gene expression. More specifically, a potential consequence of TF synergy encoded in enhancers is larger SNV effect sizes on enhancer activity. This may offer a major advantage for fine tuning gene expression levels throughout evolution. Yet, the same sensitivity also predisposes enhancers to functional disruptions: a small number of SNVs may completely abolish their activity. To buffer against such vulnerability, eukaryotic genome architectures incorporate compensatory strategies. A gene usually has multiple enhancers^53^, so failure of one can still be compensated by others. Additionally, promoters, the integration platforms for inputs from multiple enhancers, are equipped with TF redundancy and are thus robust to variability at both the DNA and TF levels. If the integration platform of promoters remains functional, other enhancers may effectively restore the regulatory output^54^. Importantly, by studying the native human genomic sequences, rather than synthetic ones, we discovered evolutionary favored motif organization rules that improve regulatory robustness, flexibility, and evolvability.

We acknowledge several current limitations of our work and highlight directions for future improvements and expansion. Firstly, the putative TFBSs analyzed here were derived from predicted TF motifs. While these are generally supported by high ISA scores, they have not been experimentally confirmed. Secondly, limited by the receptive field of DeepCompARE, we are unable to study interactions between TF motif pairs positioned beyond 255 bp from each other. Thirdly, since we only investigate naturally co-occurring TFBS pairs in native human regulatory elements, we cannot analyze the cooperativity of all TF combinations. For example, independent TFs are under-represented in DHSs (**Figure 6b**), which may have caused a weaker understanding of these by DeepCompARE, potentially leading to low ISA scores and sparse data for their cooperativity. Finally, most TF-TF PPI occupy 2-40 nm, corresponding to 6-118 nucleotides (TF proteins are 1-10 nm in size, with IDRs ranging between 1-20 nm, and the distance between two DNA nucleotides in dsDNA is on average 0.34 nm), which agrees with the observed genomic distance constraints for TF cooperativity (**Figure 3c**). Together with experimental IP-MS data, this suggests that DeepCompARE-derived cooperativity scores can provide a link between interactions at genomic and proteomic scales. However, synergy scores are inherently mutual and therefore cannot fully distinguish between different modes of TF cooperativity including 1) direct PPI, 2) indirect PPI mediated by a third partner, and 3) temporal dependency, including those established pioneer factors. Distinguishing between these modes will require temporal data or multimodal deep learning models that can reveal causal effects. We expect our framework to inform the interpretation of such future models.

## Data availability

**Supplementary Table 1: ENCODE accession numbers for external datasets used in this study**.

https://github.com/anderssonlab/DeepCompARE/blob/main/Filter_regions/encode_accession.txt

**Supplementary Table 2: TF pair synergy scores for HepG2**. https://github.com/anderssonlab/DeepCompARE/blob/main/Pd6_TF_cooperativity/tf_pair_synergy_score_hepg2_pe.csv

For column descriptions, see: https://github.com/anderssonlab/DeepCompARE/blob/main/Pd6_TF_cooperativity/table_column_explanations.txt

**Supplementary Table 3: TF pair synergy scores for K562**.

https://github.com/anderssonlab/DeepCompARE/blob/main/Pd6_TF_cooperativity/tf_pair_synergy_score_k562_pe.csv

For column descriptions, see: https://github.com/anderssonlab/DeepCompARE/blob/main/Pd6_TF_cooperativity/table_column_explanations.txt

**Supplementary Table 4: TF-level synergy scores for HepG2**.

https://github.com/anderssonlab/DeepCompARE/blob/main/Pd6_TF_cooperativity/tf_synergy_score_hepg2_pe.csv

For column descriptions, see: https://github.com/anderssonlab/DeepCompARE/blob/main/Pd6_TF_cooperativity/table_column_explanations.txt

**Supplementary Table 5: TF-level synergy scores for K562**.

https://github.com/anderssonlab/DeepCompARE/blob/main/Pd6_TF_cooperativity/tf_synergy_score_k562_pe.csv

For column descriptions, see: https://github.com/anderssonlab/DeepCompARE/blob/main/Pd6_TF_cooperativity/table_column_explanations.txt

## Code availability

https://github.com/anderssonlab/DeepCompARE

## Acknowledgements

R.A. acknowledges support from the Novo Nordisk Foundation (NNF20OC0059796) and the Novo Nordisk Foundation Center for Genomic Mechanisms of Disease (NNF21SA0072102). S.R. acknowledges support from the Novo Nordisk Foundation (NNF24SA0092867). K.L. acknowledges support from the Novo Nordisk Foundation Center for Genomic Mechanisms of Disease (NNF21SA0072102), the Novo Nordisk Foundation (NNF21CC0073729), the Stanley Center for Psychiatric Research, the US National Institute of Mental Health (R01 MH109903 and U01 MH121499), the Simons Foundation Autism Research Initiative (awards 515064 and 735604), and the Lundbeck Foundation (R223-2016-721 and R350-2020-963). R.K. acknowledges support from the Villum Foundation (1624-00072111) and the Lundbeck Foundation (R370-2021-924).

We thank Milena Rankilde Lalic and João Mendonça for help with early model interpretations. We thank Jesper Madsen, Albin Sandelin, Noam Shoresh, members of the Andersson lab, Lage lab, and the Novo Nordisk Foundation Center for Genomic Mechanisms of Disease for feedback on the DeepCompARE model and results.

## Author contributions

X.H. led the project as first author, developed the DeepCompARE model and training strategy, developed the in silico ablation approach, and performed model interpretation analyses; X.H. and R.A. conceptualized the study, formalized measures of cooperativity, and wrote the manuscript with input from all authors; H.E., S.R., R.C., R.K., and R.A. contributed to model interpretation; P.P. performed IP-MS experiments; X.H., P.P., D.H., M.R.L. and K.L. analyzed and interpreted protein-protein interaction data; C.V. and R.K. performed CAGE library preparation and sequencing; R.A. provided the overall supervision for the project.

## Conflict of interests statement

K.L. is equity holder at Sidera and consultant at Sidera and ZS Associates.

## Methods

### Genome build

All analyses and coordinates are reported using human genome reference GRCh38.

### CAGE experiment

#### Cell culture

K562 [ATCC-CCL-243] and HepG2 [ATCC-HB-8065] cells were acquired via LGC Standards. HepG2 cells were cultured in DMEM [ThermoFisher Scientific, 41965039] including 10% v/v FCS, 1% v/v Penicillin-Streptomycin [ThermoFisher Scientific, 15140122] and 1% v/v L-Glutamine [ThermoFisher Scientific, 25030081] to 75-80% confluence before trypsinization with 0.25% Trypsin-EDTA [Gibco, ThermoFisher Scientific, 25200056] and splitting 1:5-1:8. K562 cells were cultured in Advanced RPMI 1640 medium [ThermoFisher Scientific, 12663012] including 10% v/v FCS, 1% v/v Penicillin-Streptomycin and 1% v/v L-Glutamine at a concentration of 2,5-7,5×10^6 cells/ml. Both cell lines were reared in Corning T-75 flasks at 37°C with 5% CO2.

#### SLIC-CAGE sample and library preparation

HepG2 cells were washed in their culture flasks with PBS, trypsinized with 0.25% Trypsin-EDTA [Gibco, ThermoFisher Scientific, 25200056], harvested and washed once again with PBS. The K562 suspension cells were in turn washed once with PBS. Cells were counted with the Countess II FL Automated Cell Counter [ThermoFisher Scientific, AMQAF1000] and disposable cell counting chamber slides [ThermoFisher Scientific, C10228]. For both cell lines, 3 replicates of each 2,5×10^6 cells were prepared and dissolved immediately in 300 µl of RNA lysis buffer from the PureLink RNA Mini Kit [ambion, ThermoFisher Scientific, 12183025] including 1% beta-Mercaptoethanol [Sigma-Aldrich, M6250]. Due to loss of nuclei and RNA during the nuclei extraction procedure, 2×10^7 cells were prepared in triplicates for both cell lines. These nuclei samples were dissolved in 500 µl hypotonic lysis buffer (10mM Tris-HCl, ph7.5; 10mM NaCl; 3mM MgCl2; 0.1% v/v NP-40; 0.1% v/v Tween20; 1% Digitonin [Promega, G9441]), incubated for 5 min at 4°C, sheared by taking the samples up 10 times into a 1 ml syringe assembled with a 27G needle. The resulting nuclei suspension was diluted with 9,5 ml hypotonic resuspension buffer (10mM Tris-HCl, ph7.5; 10mM NaCl; 3mM MgCl2) to dilute the detergents, centrifuged for 10 min at 4°C and 500xg and the resulting nuclei pellet was dissolved in 300 µl of RNA lysis buffer from the PureLink RNA Mini Kit. RNA extraction for all nuclei and cell replicates was performed with the PureLink RNA Mini Kit according to the manufactures protocol including on-column DNA digest with the PureLink DNase Set [ThermoFisher Scientific, 12185010]. All samples were quality checked by automated gel electrophoresis of 2,5 ng per replicate with the RNA 6000 Pico kits [Agilent, 5067-1513] on a 2100 Bioanalyzer.

SLIC-CAGE libraries were prepared as outlined in detail by Cvetesic, N et al., 2018. Briefly, 2000 ng of total RNA from each nuclear or cellular replicate were mixed with 3000 ng prepared carrier RNA. First-strand cDNA synthesis was performed with these carrier-combined samples as input material. The resulting RNA:cDNA hybrids were oxidised to open the end-standing RNA nucleotides, especially the 7-methylguanosine (m7G), allowing biotinylation of capped RNA species only. Overhanging RNA stretches that are not part of the RNA:cDNA hybrids are digested with RNase ONE to ensure that the following pull-down with paramagnetic Streptavidin beads (i.e., cap-trapping) only captures the biotinylated m7Gs. cDNAs corresponding solely to capped RNAs were released from beads and hybrids by a combined RNaseH and RNase ONE digestion, the Illumina-compatible 5’ and 3’ nAnT-iCAGE linkers ligated and the second-strand cDNA synthesised. Carriers were digested, the resulting library size selected and PCR amplified. The average library length was estimated based on the fragment distribution as measured on the 2100 Bioanalyzer with the High Sensitivity DNA kit [Agilent, 2067-4626] and quantified with the Quant-iT PicoGreen dsDNA kit [ThermoFisher Scientific, P7589]. Equimolar pooling of sets of 8 samples corresponding to the full set of available adapters preceded single-end sequencing on a NextSeq550 Illumina sequencer with High Output v2.5 (75 cycles) reagents [Illumina, 20024906] using standard Illumina primers.

#### CAGE data analysis

CAGE libraries were processed through several steps to ensure high-quality data for downstream analysis. Initially, quality control of the raw sequencing reads was performed using FastQC (https://www.bioinformatics.babraham.ac.uk/projects/fastqc/) to assess the general quality. The raw FASTQ files were then trimmed using cutadapt^55^ to remove the first 5 bases from each read, eliminating adapter sequences and trim low-quality bases from the 3’ end. Reads with a length greater than 30 bases were retained for further analysis. Next, the reads were filtered using fastq_quality_filter (FASTX Toolkit, http://hannonlab.cshl.edu/fastx_toolkit/) to ensure that only high-quality reads remained. This filtering step required a minimum base quality of 30 and a passing percentage of at least 50% of the bases in each read. Following this, rRNA sequences were removed using the rRNAdust tool (https://github.com/genome/sequtils) to ensure that only relevant mRNA and other non-rRNA sequences remained for mapping. Filtered reads were mapped to the hg38 reference genome using BWA mem^56^. The resulting SAM files were then converted to BAM format using samtools^57^, followed by sorting with samtools to prepare the data for subsequent analysis. After mapping, the resulting BAM files were used to generate CTSS bed files, which were then converted into bedgraph files for the plus and minus strands and subsequently transformed into bigwig files for further analysis, using bedGraphToBigWig.

Downstream CAGE analysis was performed using PRIME (https://github.com/anderssonlab/PRIME). First, quantifyCTSSs was called on all 12 samples. The resulting object was then split by cell line using the function subsetBySupport(unexpressed=1, minSamples=0). The candidate regions were identified using the function divergentLoci. To determine the noise threshold, we selected 270,179 intergenic regions of 600bp from the human genome that don’t overlap with ENCODE representative DHS^29^. The 99.5-th percentile of CAGE expression on those intergenic regions averaged across 12 samples was set to be the noise threshold (equivalent to a CTSS count threshold = 6).

### PPI experiment

#### Protein extraction

About 500M K562 cells [ATCC-CCL-243] were collected by centrifugation (500g, 5 min, 4°C) and cell pellets were washed twice in ice-cold PBS. Cell pellet was resuspended in 18 ml of IP Lysis Buffer (Thermo Fisher Scientific), with freshly added Halt protease and phosphatase inhibitor cocktail (Thermo Fisher Scientific). After a 30 min incubation time on ice, cells were collected by centrifugation (14,000 g, 20 min, 4°C) and lysate was transferred into a clean tube. Total protein content of samples was quantified using Thermo Fisher BCA protein assay.

#### Immunoprecipitation

For each individual experiment, 2 mg of fresh protein extract were used for immunoprecipitation (IP). IP of each bait was performed in triplicate, with one triplicate of IP with isotype-matched IgG antibody used as a control. Protein lysate was incubated at 4°C overnight in the presence of 2 μg of the relevant antibody. On the next day, 90 μL of Protein A/G beads (Pierce) were added to each sample and incubated at 4°C for 4 hours. Flow-through was discarded and beads were washed once with 1 mL lysis buffer (Pierce) supplemented with Halt protease and phosphatase inhibitor cocktail (Thermo Fisher Scientific), and twice with PBS. Beads were resuspended in 100 μL of PBS and 90 µL aliquot was taken out and stored at −80°C and subsequently used for mass spectrometry analysis.

Antibodies used for immunoprecipitation were rabbit anti-MAFG (Invitrogen, PA5-30086), rabbit anti-IKZF1 (Proteintech, 12016-1-AP), rabbit anti-RFX5 (Bethyl, A300-030A), rabbit anti-RREB1 (Abcam, ab113287), rabbit anti-BACH1 (Invitrogen, PA5-117013), and rabbit anti-IgG (Abcam, ab37415).

#### Mass spectrometry for IP-MS experiments

Mass spectrometry analysis of IP samples was performed at the Taplin Mass Spectrometry Facility. IP samples on beads were washed at least five times with 100 µl 50 mM ammonium bicarbonate then 5 µl (200ng/µl) of modified sequencing-grade trypsin (Promega) was spiked in and the samples were placed in a 37ºC room overnight. The samples were then placed on a magnetic plate and the liquid removed. The extracts were then dried in a speed-vac (~1 hour). Samples were then re-suspended in 50 µl of HPLC solvent A (2.5% acetonitrile, 0.1% formic acid) and desalted by STAGE tip^58^.

On the day of analysis, the samples were reconstituted in 10 µl of HPLC solvent A. A nano-scale reverse-phase HPLC capillary column was created by packing 2.6 µm C18 spherical silica beads into a fused silica capillary (100 µm inner diameter x ~30 cm length) with a flame-drawn tip^59^. After equilibrating the column 4 µl of each sample was loaded via a Famos autosampler (LC Packings) onto the column. A gradient was formed, and peptides were eluted with increasing concentrations of solvent B (97.5% acetonitrile, 0.1% formic acid). As peptides eluted, they were subjected to electrospray ionization and then entered into an Orbitrap Fusion Lumos mass spectrometer (Thermo Scientific). Peptides were detected, isolated, and fragmented to produce a tandem mass spectrum of specific fragment ions for each peptide.

Peptide sequences (and hence protein identity) were determined by matching the UniProt human protein database (release 2023_01) with the acquired fragmentation pattern using Sequest (Thermo Scientific)^60^. The database included a reversed version of all the sequences and the data were filtered to between 1-2% peptide false discovery rate. Protein quantification was performed using GFY Core Version 3.8 (Harvard University, Cambridge, MA).

#### IP-MS data analysis

Each IP-MS dataset consisting of triplicate bait IP vs triplicate IgG control was processed as follows: (1) protein-level quantified data were log2 transformed and protein intensities were median normalized; (2) contaminants and proteins with no human gene name, detected with < 2 unique peptides, or detected in < 2 bait IP samples were removed; (3) missing values for the remaining proteins in each sample were imputed^61^ by drawing from a normal distribution with mean of μ - 1.8σ and standard deviation of 0.3σ, where μ and σ are the mean and standard deviation of the observed protein intensities, respectively; (4) limma-based^62^ two-tailed two-sample moderated t-test was performed as implemented in Genoppi^63^ (development branch v1.1.3), to calculate enrichment statistics (log2 fold change, p-value, and false discovery rate) of each protein in bait vs. control IPs; (5) proteins with log2 FC > 0 and FDR ≤ 0.1 were defined as significant interactors of the bait protein.

### Data-centric model development

DeepCompARE employed a multitasking framework, to simultaneously learn the signals of CAGE-Seq, DNase-Seq, STARR-Seq and SuRE-seq^30^ given DNA sequence. DNase-seq and STARR-Seq are downloaded from ENCODE, see **Supplementary Table 1** for accession number.

To exploit the non-orthogonality of the four data modalities, minimize label conflicts and maximize data quality, all peaks reproducible across 2 biological replicates of the same modality were included in the training set. Peaks that were irreproducible within a modality, but overlapped reproducible peaks of any other modality within the same cell type for over 80% of its length were also included.

Loose positives were defined as candidate peaks that had relatively high signal (see below), yet failed to reach significance level to be called true peaks. We observed that, compared to training on all regions including loose positives, training on a credible subset that either displayed significant signal for at least one data modality, or always being undisputedly negative for all 4 data modalities yielded better Pearson correlation on a common test set (**Supplementary Figure 1d**). Hence, loose positives were excluded from training, validation or test dataset. Below are the definitions for reproducible, irreproducible and loose positive peaks for each data modality.

For CAGE, candidate genomic regions where at least two samples of the same cell type showed expression exceeding the noise threshold were defined as reproducible. Regions, where only one sample had expression exceeding the noise threshold were defined as irreproducible. The remaining candidate regions were considered loose positives.

For DNase-seq, peaks of the same cell type overlapping at least 120 bp across replicates were considered reproducible. The remaining peaks were considered irreproducible. Due to the high quality of DNase-seq data, no peaks were considered loose positive.

For STARR-seq, peaks of the same cell type overlapping at least 400 bp across replicates were considered reproducible. The remaining peaks were considered irreproducible. To define the loose positives, STARRPeaker (version 1.0) was run on replicate-merged bam files, with parameter settings length as 500, step as 100, and FDR threshold as 0.2. All called peaks excluding the reproducible STARR-defined regulatory elements were considered loose positives.

SuRE-seq^30^ files were downloaded from https://osf.io/w5bzq/wiki/home/?view. Signals of plus and minus strand were added and analyzed with MACS3 using the command “macs3 bdgpeakcall -l 300

-g 150”. Fold-change cut-off parameter was set to 4 for true positive SuRE-seq peaks, and 2 for loose positives. Peaks overlapping with any SNPs were removed from the positive set, but were retained in the loose positive set to prevent problematic regions from getting negative labels. SuRE-seq peaks of the same cell type overlapping in at least two samples for at least 240 bp were considered reproducible. The remaining peaks were considered irreproducible. Peaks with fold change above 2 but below 4 were considered loose positives.

### Classification and regression label preparation

To define negative samples for classification, we combined ENCODE representative DHSs^29^ resized to 600bp with 600 bp human genome tiles that don’t overlap representative DHSs nor ENCODE blacklist. Then, we removed the regions overlapping at least 1 bp with positive peaks or loose positive peaks. Finally, the remaining regions were downsampled randomly to have the same number as positive peaks.

Regression prediction targets were defined as follows:

- CAGE: average Tags Per Million (TPM) across the 600 bp peak
- DNase-seq: sum of read-depth normalized signals across the 600 bp peak, averaged between replicates
- STARR-seq: sum of control-normalized signals across the 600 bp peak
- SuRE-seq: sum of fold change across the 600 bp peak

All signal values were log-transformed. We observed that classification tasks stabilized the training process of the regression model (**Supplementary Figure 1c**).

### DeepCompARE model architecture and hyperparameter selection

DeepCompARE employs a multitasking framework that takes as input a one-hot encoded DNA sequence of 600bp (A: [0,0,0,1], C: [0,1,0,0], G: [0,0,1,0], T: [0,0,0,1], N: [0,0,0,0]). The output for one sequence is a vector of 16 values: predicted average signal across the sequence measured by CAGE (HepG2), CAGE (K562), DNase-seq (HepG2), DNase-seq (K562), STARR-seq (HepG2), STARR-seq (K562), SuRE-seq (HepG2), SuRE-seq (K562) respectively, followed by 8 binary classification values in identical order.

Inspired by BPNet^22,64^, we used exponentially dilated convolution to achieve large receptive fields in deep layers. We also avoided pooling to maintain base pair resolution during backtracking for feature importance. DeepCompARE used only 5 convolutional layers without residual connections followed by only one fully connected layer. We adjusted the kernel widths to [15,9,9,9,9] and the number of channels in each layer to 128. The dilation rate was [1,2,4,8,16]. To remove the unnecessary connections and interference between channels calculating different tasks, we set the “group” parameter in torch.nn.Conv1d function to 4 and 8 for the 4th and 5th layers respectively, which reduced the total number of parameters by approximately half. Absence of grouping in the first 3 layers allows the model to learn sequence features shared by all assays.

To avoid potential imbalance between regression tasks due to different scale of measurement, the weights for tracks were [1,1,8,8,5,5,1,1].

The learning rate was fixed at 0.001. Batch size was set to 4096. Dropout layer with dropout rate 0.1 was applied after each convolutional layer to reduce overfitting. Reverse complement data augmentation was applied to further decrease overfitting.

### SNV effect size prediction and benchmarking against Enformer

Given the locus of an SNV, we extended the region to 600 bp with the SNV fixed at center and input the reference and alternative sequence to each model. The difference in model output was considered the effect size of the SNV. Thus, DeepCompARE made 4 predictions from regression tracks for CAGE, DNase-seq, STARR-seq and SuRE-seq tracks in matched cell types (for HEL 92.1.7 data, we used K562 tracks). For Enformer, we calculated changes in CAGE and DNase-seq predictions, in both a cell-type matched and cell-type agnostic manner.

### Identification of promoters and enhancers

Promoters were defined to be CAGE peaks within 500 bp of annotated gene TSSs, as defined by UCSC knownGene. Enhancers were defined to be the overlapping peaks of DNase-seq and STARR-seq, regardless of genomic location.

### Single base, TFBS, and TF-level ISA scores

A single base ISA score was calculated as the difference in model output caused by replacing the given base by “N”, encoded as [0,0,0,0], consistent with the encoding for “unknown base” during training. Bases whose perturbation reduced the model output were considered positive importance scores.

For each putative TFBS in the JASPAR database^36^ with matching score exceeding 500, we ablated the motif by replacing all its nucleotides to “N” while leaving the rest of the sequence intact. The difference in model output caused by the ablation was defined as the corresponding motif importance. TFBSs whose ablation reduced the model output were considered positive importance scores.

Since TF effect varies with sequence context, to obtain an overall assessment of the effect of a TF to a certain regulatory activity (TF-level ISA), we aggregated the importance scores of all its TFBSs across all considered sequences. We used average motif importance as a basic aggregation method whenever the importance scores were directly comparable. When TF importances were derived from different tracks and thus are not directly comparable, we used the Kolmogorov-Smirnov test to calculate the difference in cumulative distributions between the importances of a target TF and the importances of all TFs assigned by the same track, then used the signed D-statistics as the normalized aggregated importance score. Specifically, if the median of the importance of a target TF was smaller than the overall median, we negated the D-statistics to indicate an importance below median.

For gnomAD, phyloP and TF cooperativity analysis, we further required each considered TF gene to be expressed above 0.5 transcripts per million (TPM), to increase the likelihood that the TF is functional in the corresponding cell type.

### Enrichment analysis of gnomAD non-rare variants

For all loss-of-function SNVs in gnomAD that passed all quality filters, we selected those overlapping promoters or enhancers of HepG2 or K562, binned the SNVs by predicted effect size, and calculated the odds ratio of seeing a non-rare allele within each bin. Non-rare alleles were defined by several allele frequency thresholds (0.0001, 0.001, 0.01, 0.05) to ensure robustness of the analysis.

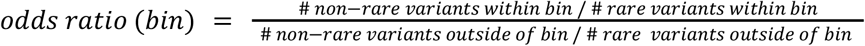

### Formal definition of cooperativity: redundancy and synergy

We defined cooperativity using a top-down approach, i.e., a positive motif effect indicates a decrease of sequence activity upon perturbation. Let the predicted activity for the original intact sequence be a, the output for the sequence with TFBS_A_ ablated be a_-A_, the output for sequence with TFBS_B_ ablated be a_-B_, and the output for sequence with both motifs ablated be a_-AB_. Assume the effects of TFBS_A_ and TFBS_B_ in the intact sequence are e_A_ and e_B_, and the combinatorial effect of the (TFBS_A_, TFBS_B_) pair is e_AB_, then we have:

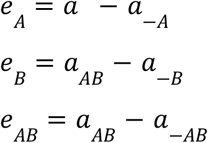

Let e_A|-B_ be the effect of TFBS_A_ without TFBS_B_, e_B|-A_ be the effect of TFBS_B_ without TFBS_A._ Then, by definition, the combinatorial effect of losing both TFBSs (e_AB_) is the same as the effect of losing TFBS_A_ first (*e*_A_), then losing TFBS given that TFBS_A_ is already lost (*e*_*A B*|−*A*_). That is:

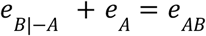

Similarly:

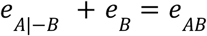

If TFBS_A_ and TFBS_B_ contribute independently to the overall activity, then the effect of TFBS_B_ should not be affected by presence or absence of TFBS_A_.

That is, under **independence**:

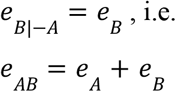

Thus, independence implies additivity of TFBS effects measured by ablation under a logarithmic scale.

**Redundancy** is defined as:

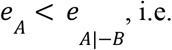

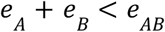

**Synergy** is defined as:

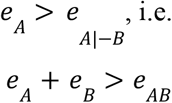

Note that both synergy and redundancy are mutual by definition. If we know that TFBS_A_ depends on TFBS_B_ to achieve full function, then TFBS_A_ depends on TFBS_B_ to the same degree.

In principle, cooperativity can also be defined using a bottom-up approach (**Supplementary Figure 6d**). A TF effect in the bottom-up approach is implicitly conditioned on the absence of a partner.

### Relationship between redundancy and subadditivity

In our definition, redundancy is *e*_*AB*_ > *e*_*A*_ + *e*_*B*_, meaning that the combinatorial effect is larger than the sum of individual effects measured top-down (in intact sequence context), i.e., superadditivity. At a first glance, our definition appears to contradict the mainstream notion that redundancy implies subadditivity. However, a more nuanced analysis using the above mathematical definitions that explicitly accounts for the presence or absence of the TFBS partner resolves this paradox.

Suppose TFBS_A_ and TFBS_B_ are redundant, meaning *e*_*B*_ < *e*_*B*|−*A*_. Then *e*_*A*|−*B*_ + *e*_*B*_ < *e*_*A*|−*B*_ + *e*, i.e. *e*_*AB*_ < *e*_*A*|−*B*_ + *e*_*B*|−*A*_, meaning the combinatorial effect is smaller than the sum of individual effects measured bottom-up (in background sequence), in line with the claim that redundancy leads to subadditivity. The same analysis applies to synergy and superadditivity.

### Independence, synergy and redundancy scores of TF pairs

We define interaction strength (*i*) between TFBS_A_ and TFBS_B_:

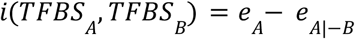

which implies that:

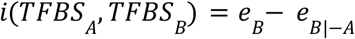

Hence, a TF pair has one unique cooperativity value given a sequence context, where a positive cooperativity indicates synergy (both TFs need each other to achieve full functionality) and negative cooperativity indicates functional redundancy between the pair (damage to one can be compensated by the other to various degrees).

To aggregate cooperativity over all sequence contexts for each TF pair, we define independence, synergy and redundancy scores:

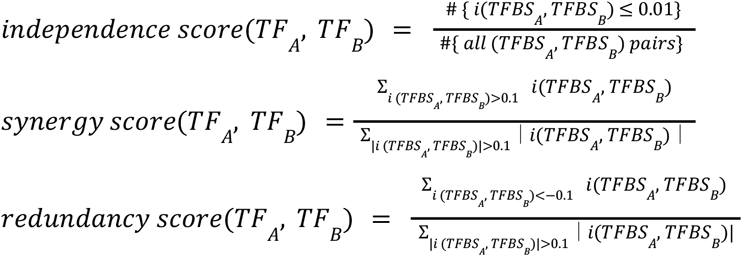

The interaction threshold ±0.1 ensures that only TFBS pairs with substantial synergy or redundancy are taken into account, in order to reduce the influence of potential noise in model predictions. Hence, the synergy (redundancy) score is simply the fraction of substantially synergistic (redundant) cases among all cooperative cases, weighted by interaction strength. Then, by definition,

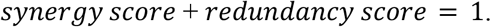

With interaction threshold ±0.1, the four modalities almost always agree on the cooperativity of all TF pairs, suggesting that cooperativity of a TF pair is robust to the functional assays once the sequence context is fixed. The <2% TF pairs with inconsistent predictions between tracks were excluded from downstream analysis. Since only activators, not repressors, were of interest in this study, we also removed the TF pairs where either one of *e*_*A*_, *e*_*B*_, *e*_*A*|−*B*_, *e*_*B*|−*A*_ were negative. For the TF pairs with agreement across modalities, we averaged the interaction values obtained from all tracks as the final interaction value given the current sequence context.

To avoid inaccurate scores due to underrepresentation, TF pairs with fewer than 11 occurrences were excluded from independence, synergy and redundancy score calculations. TF pairs with a sum of absolute interaction lower than 1 were excluded from synergy and redundancy score calculations. As a consequence, independent TF pairs don’t have synergy or redundancy scores.

We assigned a cooperativity type to all TF pairs according to the following criteria:

- Redundant: sum of absolute interaction ≥ 1, and redundant score > 0.7.
- Synergistic: sum of absolute interaction ≥ 1, and synergistic score > 0.7.
- Intermediate: sum of absolute interaction ≥ 1, and 0.3 ≤ synergistic score ≤ 0.7.
- Independent: sum of absolute interaction < 1, and independence score > 0.9.

### TF-level synergy and redundancy scores

The synergy and redundancy scores of TF_A_ aggregates the interaction strength of all TFBS pairs containing TFBS_A_:

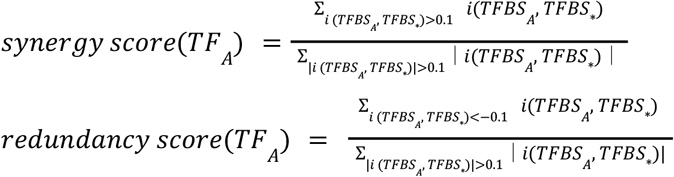

“*” denotes any TF whose binding sites co-occur with TFBS_A_ on regulatory elements. Again, by definition:

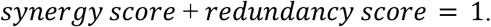

We assigned a cooperativity type to all TFs according to the following criteria:

- Redundant: sum of absolute interaction ≥ 5, and redundant score > 0.7.
- Synergistic: sum of absolute interaction ≥ 5, and synergistic score > 0.7.
- Intermediate: sum of absolute interaction ≥ 5, and 0.3 ≤ synergistic score ≤ 0.7.
- Independent: sum of absolute interaction < 5, and independence score > 0.95.

### gnomAD constraint repurposed for TFBSs

For each TF, we obtained all its putative binding site sequences across promoter-proximal or promoters-distal regions, and compared the expected (*Exp*) and observed (*Obs*) number of rare variants (minor allele frequency < 0.001). The expected number of rare variants was calculated using the trinucleotide context-specific mutation rates estimated from the gnomAD database. Then we calculated a repurposed Gnocchi^38^ constraint score for TFBSs:

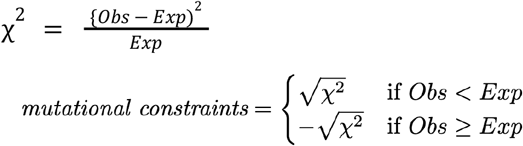

## Notes

https://github.com/anderssonlab/DeepCompARE

## References

1. Lagha, M., Bothma, J. P. & Levine, M. Mechanisms of transcriptional precision in animal development. Trends Genet. 28, 409–416 (2012).

2. Cvetesic, N. & Lenhard, B. Core promoters across the genome. Nat. Biotechnol. 35, 123–124 (2017).

3. Haberle, V. & Stark, A. Eukaryotic core promoters and the functional basis of transcription initiation. Nat. Rev. Mol. Cell Biol. 19, 621–637 (2018).

4. Andersson, R. & Sandelin, A. Determinants of enhancer and promoter activities of regulatory elements. Nat. Rev. Genet. 21, 71–87 (2020).

5. Lambert, S. A. et al. The Human Transcription Factors. Cell 172, 650–665 (2018).

6. Spitz, F. & Furlong, E. E. M. Transcription factors: from enhancer binding to developmental control. Nat. Rev. Genet. 13, 613–626 (2012).

7. Iwafuchi-Doi, M. & Zaret, K. S. Pioneer transcription factors in cell reprogramming. Genes Dev. 28, 2679–2692 (2014).

8. Thanos, D. & Maniatis, T. Virus induction of human IFN beta gene expression requires the assembly of an enhanceosome. Cell 83, 1091–1100 (1995).

9. Panne, D., Maniatis, T. & Harrison, S. C. An atomic model of the interferon-beta enhanceosome. Cell 129, 1111–1123 (2007).

10. Kulkarni, M. M. & Arnosti, D. N. Information display by transcriptional enhancers. Development 130, 6569–6575 (2003).

11. Arnosti, D. N. & Kulkarni, M. M. Transcriptional enhancers: Intelligent enhanceosomes or flexible billboards? J. Cell. Biochem. 94, 890–898 (2005).

12. Junion, G. et al. A Transcription Factor Collective Defines Cardiac Cell Fate and Reflects Lineage History. Cell 148, 473–486 (2012).

13. Kribelbauer-Swietek, J. F. et al. Context transcription factors establish cooperative environments and mediate enhancer communication. Nat. Genet. 56, 2199–2212 (2024).

14. Morgunova, E. & Taipale, J. Structural perspective of cooperative transcription factor binding. Curr. Opin. Struct. Biol. 47, 1–8 (2017).

15. De Val, S. et al. Combinatorial regulation of endothelial gene expression by ets and forkhead transcription factors. Cell 135, 1053–1064 (2008).

16. Spivakov, M. Spurious transcription factor binding: non-functional or genetically redundant? Bioessays 36, 798–806 (2014).

17. Joshua L Payne, A. W. Mechanisms of mutational robustness in transcriptional regulation. Front. Genet. 6, 378 (2015).

18. Wu, W.-S. & Lai, F.-J. Functional redundancy of transcription factors explains why most binding targets of a transcription factor are not affected when the transcription factor is knocked out. BMC Syst. Biol. 9 Suppl 6, S2 (2015).

19. Einarsson, H. et al. Promoter sequence and architecture determine expression variability and confer robustness to genetic variants. Elife 11, e80943 (2022).

20. Hollenhorst, P. C., Shah, A. A., Hopkins, C. & Graves, B. J. Genome-wide analyses reveal properties of redundant and specific promoter occupancy within the ETS gene family. Genes Dev. 21, 1882–1894 (2007).

21. Zhao, Y. et al. ‘Stripe’ transcription factors provide accessibility to co-binding partners in mammalian genomes. Mol. Cell 82, 3398–3411.e11 (2022).

22. Avsec, Ž. et al. Base-resolution models of transcription-factor binding reveal soft motif syntax | Nature Genetics. Nat. Genet. 53, 354–366 (2021).

23. de Almeida, B. P., Reiter, F., Pagani, M. & Stark, A. DeepSTARR predicts enhancer activity from DNA sequence and enables the de novo design of synthetic enhancers. Nat. Genet. 54, 613–624 (2022).

24. Sahu, B. et al. Sequence determinants of human gene regulatory elements. Nat. Genet. 54, 283–294 (2022).

25. Kim, D. S. et al. The dynamic, combinatorial cis-regulatory lexicon of epidermal differentiation. Nat. Genet. 53, 2020.10.16.342857 (2020).

26. Martyn, G. E. et al. Rewriting regulatory DNA to dissect and reprogram gene expression. bioRxiv (2023) doi:10.1101/2023.12.20.572268.

27. Kelley, D. R., Snoek, J. & Rinn, J. L. Basset: learning the regulatory code of the accessible genome with deep convolutional neural networks. Genome Res. 26, 990–999 (2016).

28. Toneyan, S. & Koo, P. K. Interpreting Cis-regulatory interactions from large-scale deep neural networks for genomics. bioRxivorg (2024) doi:10.1101/2023.07.03.547592.

29. Moore, J. E. et al. Expanded encyclopaedias of DNA elements in the human and mouse genomes. Nature 583, 699–710 (2020).

30. van Arensbergen, J. et al. High-throughput identification of human SNPs affecting regulatory element activity. Nat. Genet. 51, 1160–1169 (2019).

31. Avsec, Ž. et al. Effective gene expression prediction from sequence by integrating long-range interactions. Nat. Methods 18, 1196–1203 (2021).

32. Kircher, M. et al. Saturation mutagenesis of twenty disease-associated regulatory elements at single base-pair resolution. Nat. Commun. 10, 3583 (2019).

33. Nair, S., Shrikumar, A., Schreiber, J. & Kundaje, A. fastISM: performant in silico saturation mutagenesis for convolutional neural networks. Bioinformatics 38, 2397–2403 (2022).

34. Salvatore, M., Horlacher, M., Marsico, A., Winther, O. & Andersson, R. Transfer learning identifies sequence determinants of cell-type specific regulatory element accessibility. NAR Genom Bioinform 5, lqad026 (2023).

35. Musunuru, K. et al. From noncoding variant to phenotype via SORT1 at the 1p13 cholesterol locus. Nature 466, 714–719 (2010).

36. Rauluseviciute, I. et al. JASPAR 2024: 20th anniversary of the open-access database of transcription factor binding profiles. Nucleic Acids Res. 52, D174–D182 (2024).

37. Andersson, R. et al. Nuclear stability and transcriptional directionality separate functionally distinct RNA species. Nat. Commun. 5, 5336 (2014).

38. Chen, S. et al. A genomic mutational constraint map using variation in 76,156 human genomes. Nature 625, 92–100 (2024).

39. THE GTEX CONSORTIUM. The GTEx Consortium atlas of genetic regulatory effects across human tissues. Science 369, 1318–1330 (2020).

40. Landrum, M. J. et al. ClinVar: public archive of relationships among sequence variation and human phenotype. Nucleic Acids Res. 42, D980–5 (2014).

41. Kanai, M. et al. Insights from complex trait fine-mapping across diverse populations. Preprint at 10.1101/2021.09.03.21262975 (2021).

42. Zoonomia Consortium. A comparative genomics multitool for scientific discovery and conservation. Nature 587, 240–245 (2020).

43. Szklarczyk, D. et al. The STRING database in 2023: protein-protein association networks and functional enrichment analyses for any sequenced genome of interest. Nucleic Acids Res. 51, D638–D646 (2023).

44. Li, T. et al. A scored human protein–protein interaction network to catalyze genomic interpretation. Nat. Methods 14, 61–64 (2017).

45. Ilsley, M. D. et al. Krüppel-like factors compete for promoters and enhancers to fine-tune transcription. Nucleic Acids Res. 45, 6572–6588 (2017).

46. Hsieh, P. N., Fan, L., Sweet, D. R. & Jain, M. K. The Krüppel-like factors and control of energy homeostasis. Endocr. Rev. 40, 137–152 (2019).

47. Peng, Y. et al. Detection of new pioneer transcription factors as cell-type specific nucleosome binders. (2023) doi:10.7554/elife.88936.1.

48. Kundaje, A. et al. Integrative analysis of 111 reference human epigenomes. Nature 518, 317–330 (2015).

49. Bergman, D. T. et al. Compatibility rules of human enhancer and promoter sequences. Nature 607, 176–184 (2022).

50. Kentepozidou, E. et al. Clustered CTCF binding is an evolutionary mechanism to maintain topologically associating domains. Genome Biol. 21, 5 (2020).

51. Farré, D., Bellora, N., Mularoni, L., Messeguer, X. & Albà, M. M. Housekeeping genes tend to show reduced upstream sequence conservation. Genome Biol. 8, R140 (2007).

52. Farley, E. K. et al. Suboptimization of developmental enhancers. Science 350, 325–328 (2015).

53. Sheth, M. U. et al. Mapping enhancer-gene regulatory interactions from single-cell data. bioRxivorg 2024.11. 23.624931 (2024) doi:10.1101/2024.11.23.624931.

54. Perry, M. W., Boettiger, A. N., Bothma, J. P. & Levine, M. Shadow enhancers foster robustness of Drosophila gastrulation. Curr. Biol. 20, 1562–1567 (2010).

55. Martin, M. Cutadapt removes adapter sequences from high-throughput sequencing reads. EMBnet.journal 17, 10 (2011).

56. Li, H. & Durbin, R. Fast and accurate long-read alignment with Burrows-Wheeler transform. Bioinformatics 26, 589–595 (2010).

57. Li, H. et al. The Sequence Alignment/Map format and SAMtools. Bioinformatics 25, 2078–2079 (2009).

58. Rappsilber, J., Ishihama, Y. & Mann, M. Stop and go extraction tips for matrix-assisted laser desorption/ionization, nanoelectrospray, and LC/MS sample pretreatment in proteomics. Anal. Chem. 75, 663–670 (2003).

59. Peng, J. & Gygi, S. P. Proteomics: the move to mixtures. J. Mass Spectrom. 36, 1083–1091 (2001).

60. Eng, J. K., McCormack, A. L. & Yates, J. R. An approach to correlate tandem mass spectral data of peptides with amino acid sequences in a protein database. J. Am. Soc. Mass Spectrom. 5, 976–989 (1994).

61. Tyanova, S. et al. The Perseus computational platform for comprehensive analysis of (prote)omics data. Nat. Methods 13, 731–740 (2016).

62. Ritchie, M. E. et al. limma powers differential expression analyses for RNA-sequencing and microarray studies. Nucleic Acids Res. 43, e47 (2015).

63. Pintacuda, G. et al. Genoppi is an open-source software for robust and standardized integration of proteomic and genetic data. Nat. Commun. 12, 2580 (2021).

64. Trevino, A. E. et al. Chromatin and gene-regulatory dynamics of the developing human cerebral cortex at single-cell resolution. Cell 184, 5053–5069.e23 (2021).

