## Supplementary Material for "Genome-wide rules of transcription factor cooperativity revealed through *in silico* binding site ablation"

### Contents

- Supplementary Figures 1-10
- Supplementary Tables 1-5

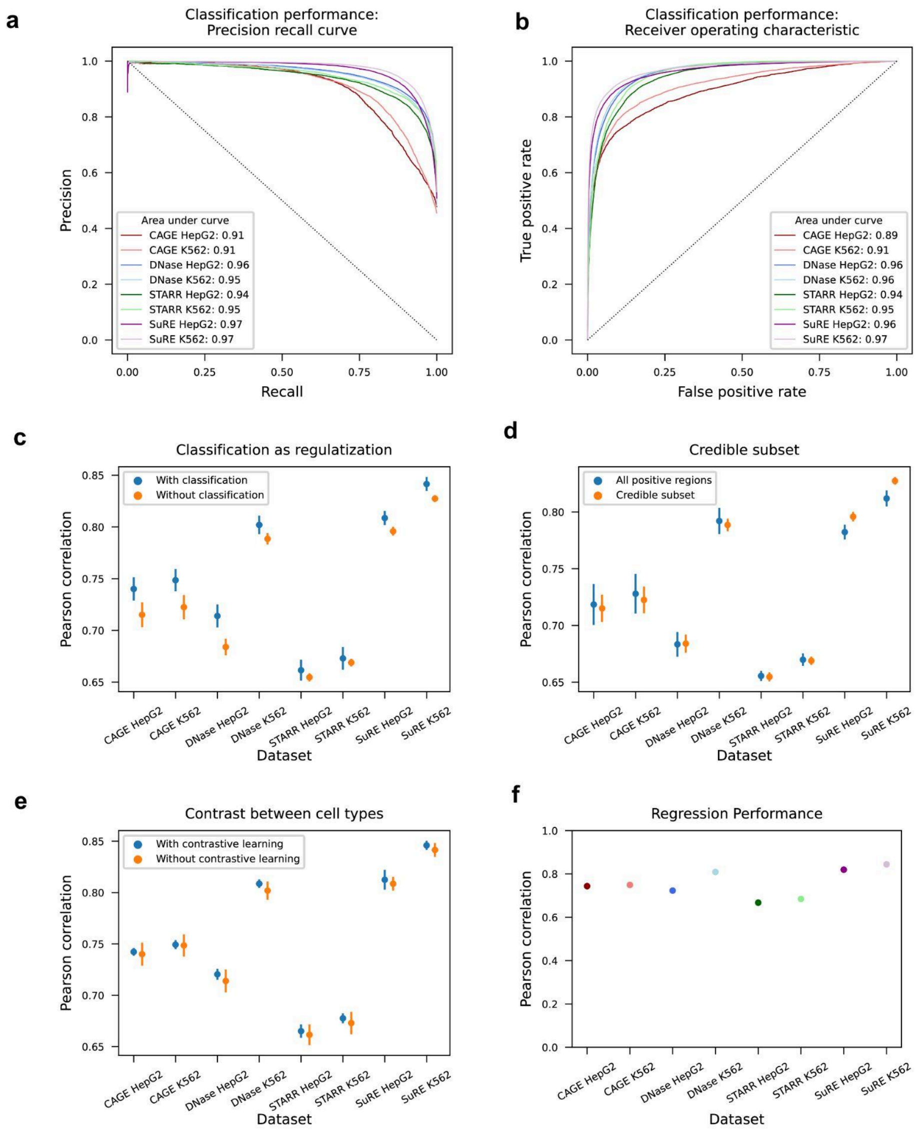


**Supplementary Figure 1. Training DeepCompARE with data-dependent regularization.**

**a:** Classification performance of DeepCompARE measured by precision recall curve. **b:** Classification performance of DeepCompARE measured by receiver operating characteristic curve. **c:** Classification as regularization improves Pearson correlation between prediction and true signal for most tracks. Error bar: 95% CI, n=5. **d:** Regression model trained on all regions with signal does not outperform regression model trained on the credible subset (Methods). Error bar: 95% CI, n=5. **e:** Contrast between cell types improves Pearson correlation. Error bar: 95% CI, n=5. **f:** Pearson correlation of prediction achieves 0.74, 0.75, 0.72, 0.81, 0.67, 0.68, 0.82, 0.84 for the final published model

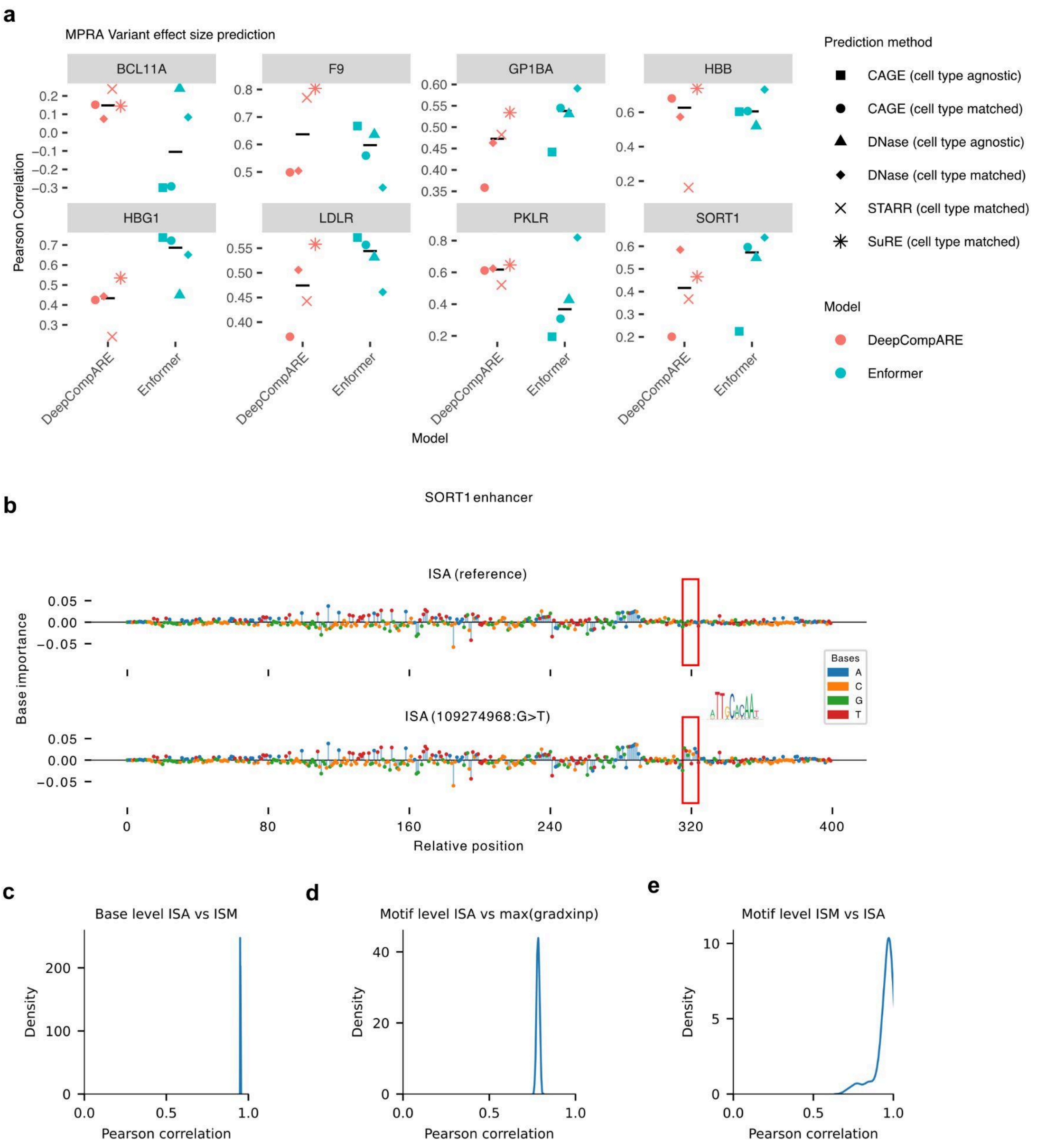


**Supplementary Figure 2. Accurate prediction of SNV effect size and efficient approximation by in silico ablation (ISA). a:** SNV effect size prediction performance of DeepCompARE and Enformer on saturation mutagenesis datasets of 8 disease-related promoters and enhancers. **b:** ISA-derived base importance scores are elevated across the de novo C/EBP binding site created by mutation chr1:109274968 G>T. **c:** Distribution of Pearson correlation between base level ISA versus ISM. 10,000 sequences were randomly sampled without replacement and flattened for correlation calculation, until all 55,077 sequences x 16 tracks = 881,232 importances scores of all promoters and enhancers of HepG2 and K562 were sampled. **d:** Distribution of Pearson correlation between motif ISA and maximum gradient x input across the motif. 100,00 motifs were randomly sampled without replacement for correlation calculation, until all the 916372 motifs were sampled. **e**: Distribution of Pearson correlation between motif ISM and ISA across the motif. Due to limitation in computation resources, 100 motifs were randomly sampled without replacement for correlation calculation, until all the 6163 motifs were sampled.

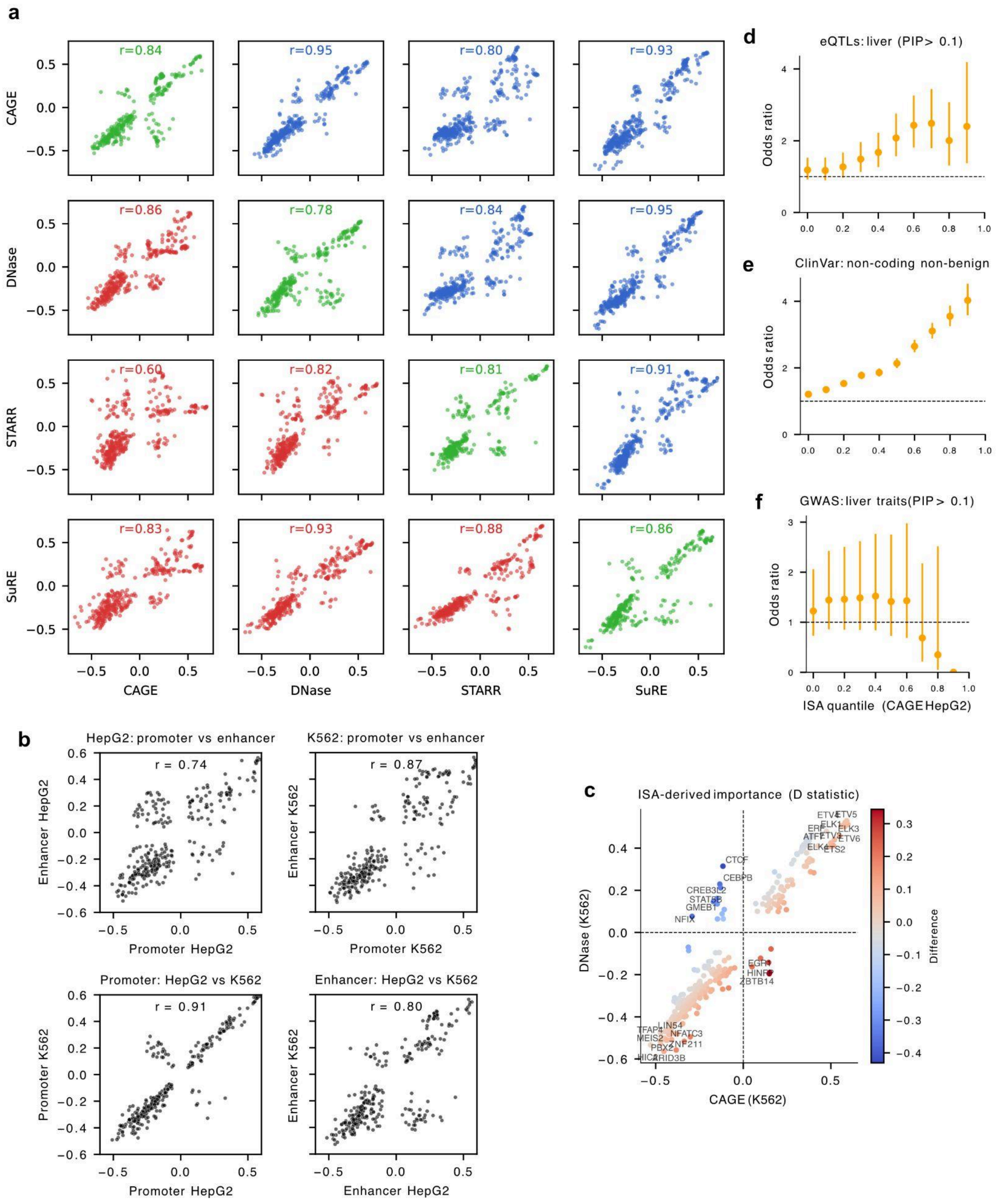


**Supplementary Figure 3. Comparison and contrast of motif ISA scores between tracks and regulatory elements. a:** Scatterplots comparing ISA-derived TF motif importance of CAGE, DNase-seq, STARR-seq and SuRE-seq tracks, as measured by signed Kolmogorov-Smirnov test D statistics (Methods). Upper diagonal (blue) depicts K562. Lower diagonal (red) depicts HepG2. Diagonal compares the motif importance derived from the same track for HepG2 (horizontal axis) and K562 (vertical axis). **b:** Scatterplots comparing ISA-derived TF motif importance aggregated from HepG2 promoters, HepG2 enhancers, K562 promoters and K562 enhancers (Methods), using CAGE track. **c:** Differences in ISA-derived TF motif importance for CAGE track (horizontal axis) versus DNase-seq track (vertical axis) in K562 cells. **d-f:** Enrichment of (**d**) fine-mapped liver eQTLs (PIP > 0.1), (**e**) ClinVar non-coding and non-benign variants, and (**f**) fine-mapped GWAS variants for liver traits (PIP > 0.1), versus common variants.

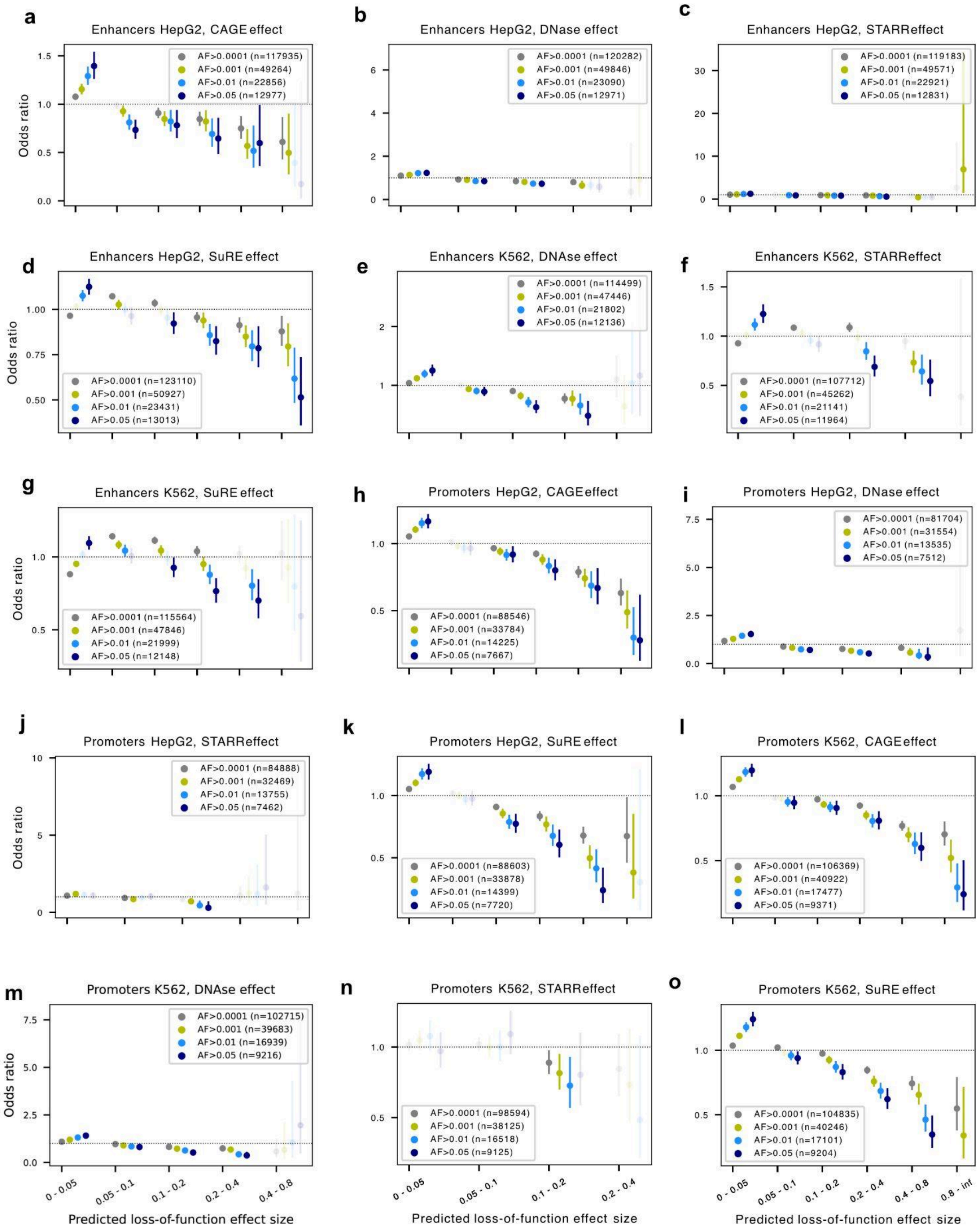


**Supplementary Figure 4. Common variant depletion of large effect size SNVs. a-o**: Predicted loss-of-function effect size on HepG2 and K562 from CAGE, DNase-seq, STARR-seq, and SuRE-seq tracks (horizontal axis), versus enrichment (odds ratio, vertical axis) of gnomAD variants, thresholded by minor allele frequency (MAF). Transparent dots represent P >= 0.05. Solid dots represent P < 0.05. Error bar: 95% CI. CAGE and SuRE-seq tracks reflect mutational constraints for both promoters and enhancers, whereas STARR-seq tracks only reflect those of enhancers.

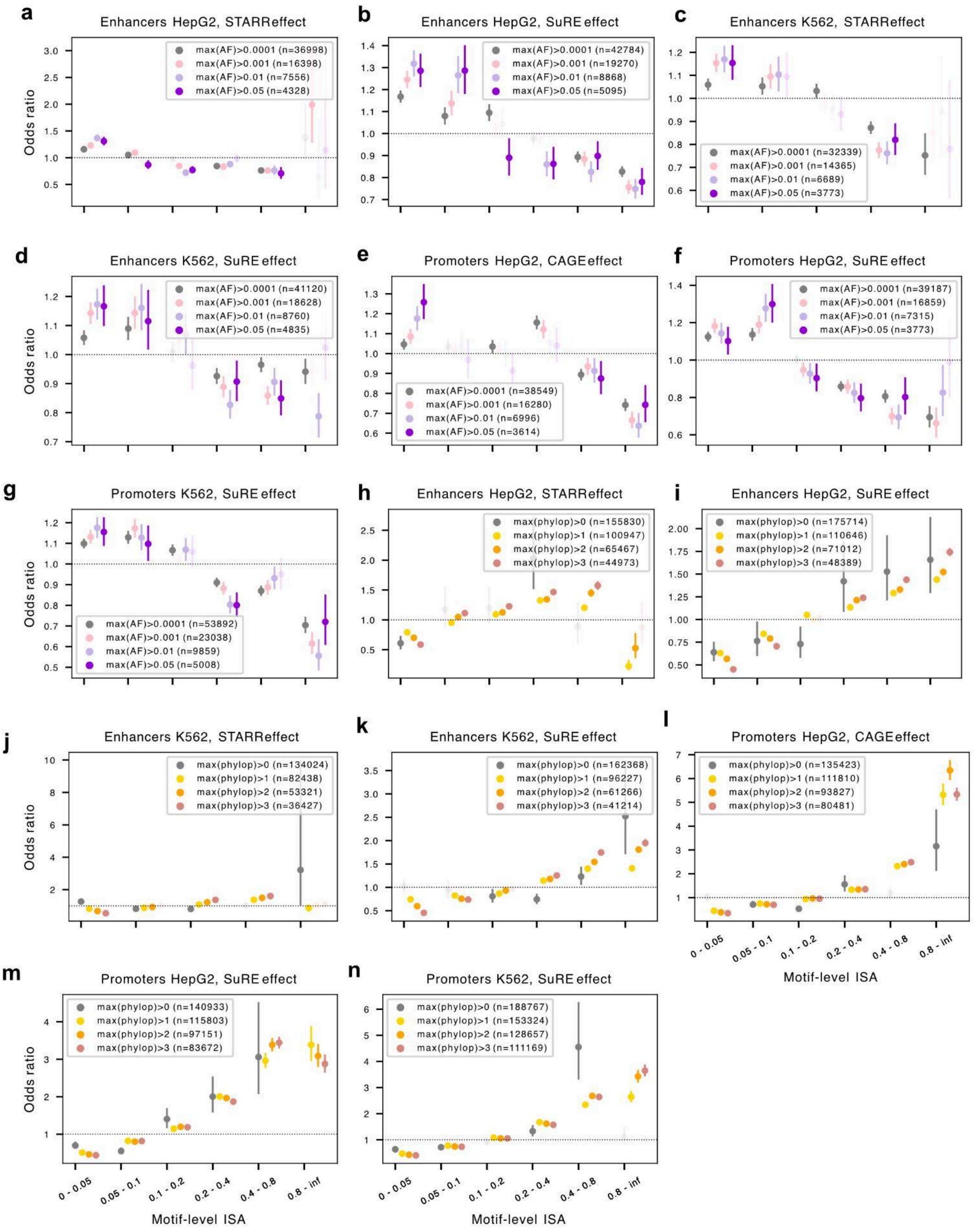


**Supplementary Figure 5. TF binding sites with large ISA scores are more conserved and less likely to harbor common variants. a-g**: Motif-level ISA scores (horizontal axis) for HepG2 and K562, versus the enrichment of gnomAD variants (odds ratio, vertical axis), thresholded by the maximum MAF across the motif. CAGE and SuRE-seq tracks are used for promoters. STARR-seq and SuRE-seq tracks are used for enhancers. Transparent dots represent P >= 0.05. Solid dots represent P < 0.05. Error bar: 95% CI. **h-n**: As **a-g**, but considering motif-level ISA scores (horizontal axis), versus enrichment (odds ratio, vertical axis) of the maximum phyloP score across the motif.

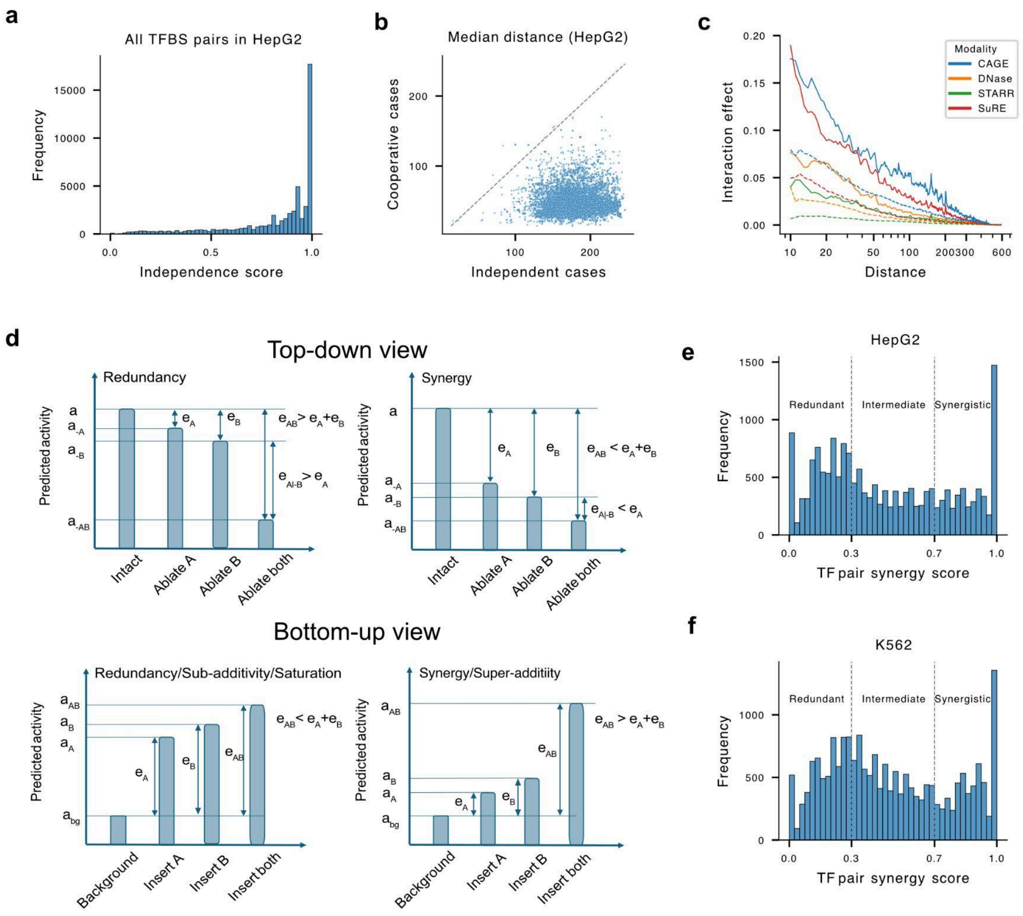


**Supplementary Figure 6. TF pair cooperativity.** **a**: Histogram of independence score for all TFBS pairs in HepG2. **b:** Median distance between independent (horizontal axis) and cooperative (vertical axis) TFBS pair instances for each TF pair. **c:** As distance increases, TF partner effect ($|e_A - e_{A|-B}|$) measured by a 6-layered CNN with a 511 bp receptive field decreases to zero. **d:** Illustration of redundancy and synergy from both top-down (upper panel) and bottom-up view (lower panel) for comparison. **e:** Histogram of TF pair synergy scores for HepG2. Gray vertical lines mark delineation of redundant (synergy score < 0.3), intermediate (0.3 ≤ synergy score ≤ 0.7), and synergistic (synergy score > 0.7) TF pairs. **f:** Same as **e**, but for K562

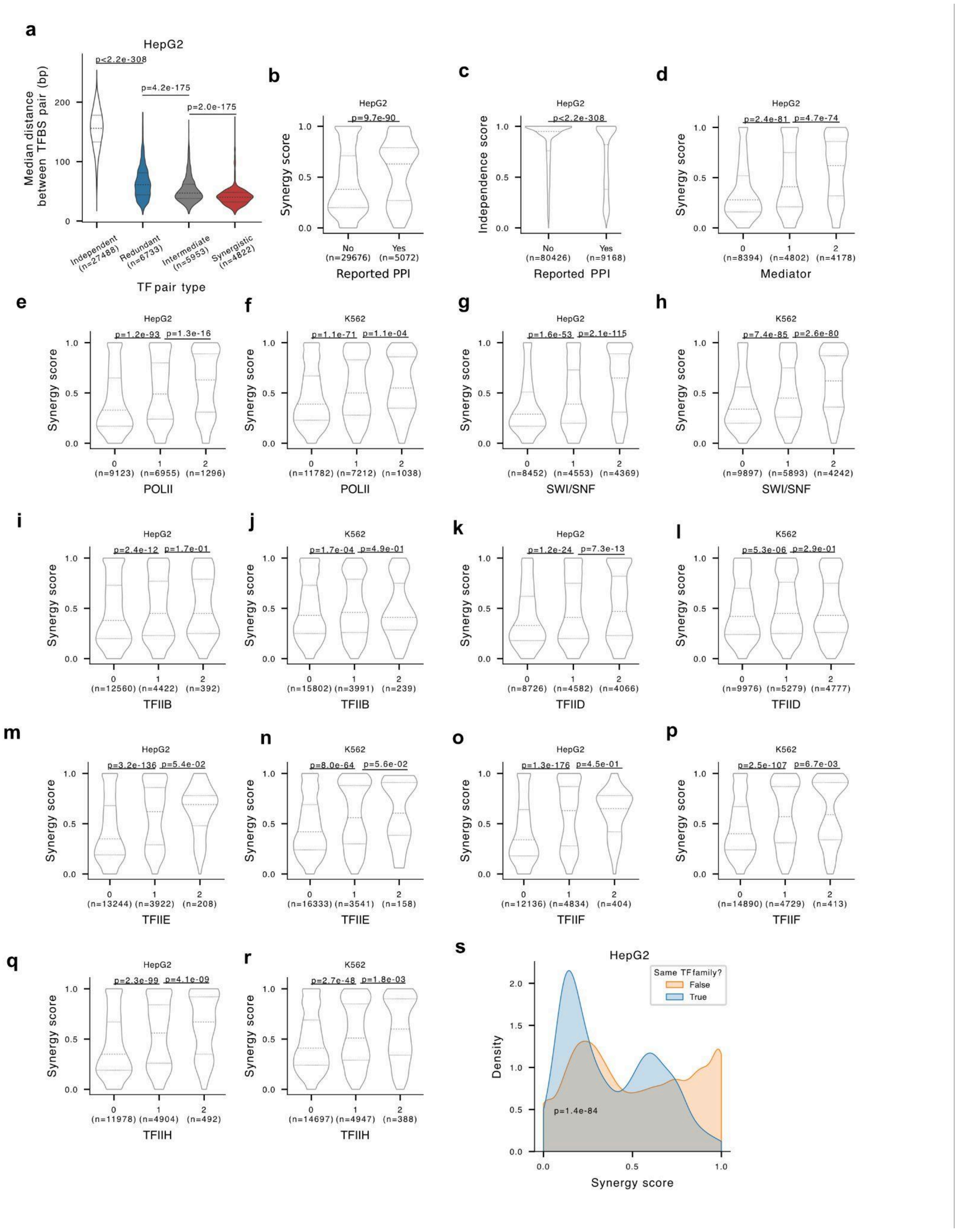


**Supplementary Figure 7. Synergy score and independence score imply direct and indirect protein-protein interaction regardless of cell types. a:** Distribution of median genomic distance between TFBS pairs for independent, redundant, intermediate, and synergistic TF pairs in HepG2. **b:** TF pairs with protein-protein interaction in public databases (STRING and InWeb) show higher synergy scores in HepG2. **c:** TF pairs with protein-protein interaction in public databases show lower independence score in HepG2. **d:** TF pairs with more Mediator complex interactors tend to have higher synergy scores in HepG2. **e-r**: TF pairs that interact with a common third factor (except TFIIB) tend to have higher synergy scores, for both HepG2 and K562, indicating that synergy may be mediated by certain common third factors. All p-values were derived from a two-sided Mann-Whitney U test. **s**: Distribution of TF pair synergy scores in HepG2, grouped by whether the TF pair belong to the same TF family or not.

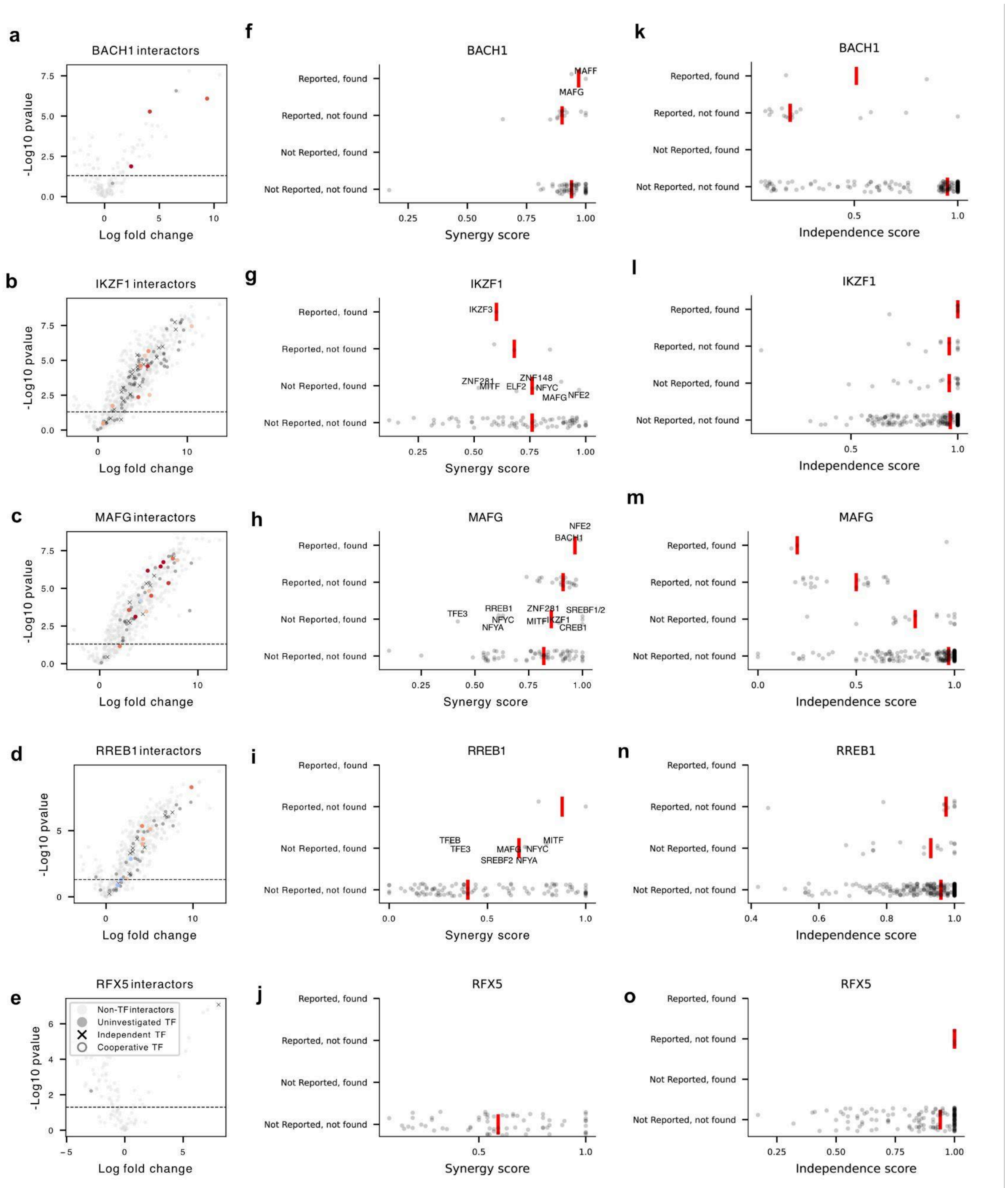


**Supplementary Figure 8. IP-MS experiment. a-e:** Volcano plot for each TF bait, with log fold change of TF bait over IgG as horizontal axis, -log10 P value from two-tailed, two-sample moderated t-test as vertical axis (Methods). Light gray dots: Protein interactors that are not TFs. Dark gray dots: TFs uninvestigated by combinatorial ISA due to lack of high-quality predicted binding site locations. Black crosses: TF partners predicted to act independently from the TF bait. Colorful dots: Cooperative TFs, colored by synergy score. **f-j:** Synergy score distribution of all investigated TF partners of each TF bait, grouped by whether their interactions are reported by public databases, and whether they are found by IP-MS experiment. Red vertical line indicates the median synergy score within each group. TFs found in the IP experiment have their names annotated at corresponding groups, and corresponding synergy scores. **k-o:** Same as **f-j**, but considering independence score.

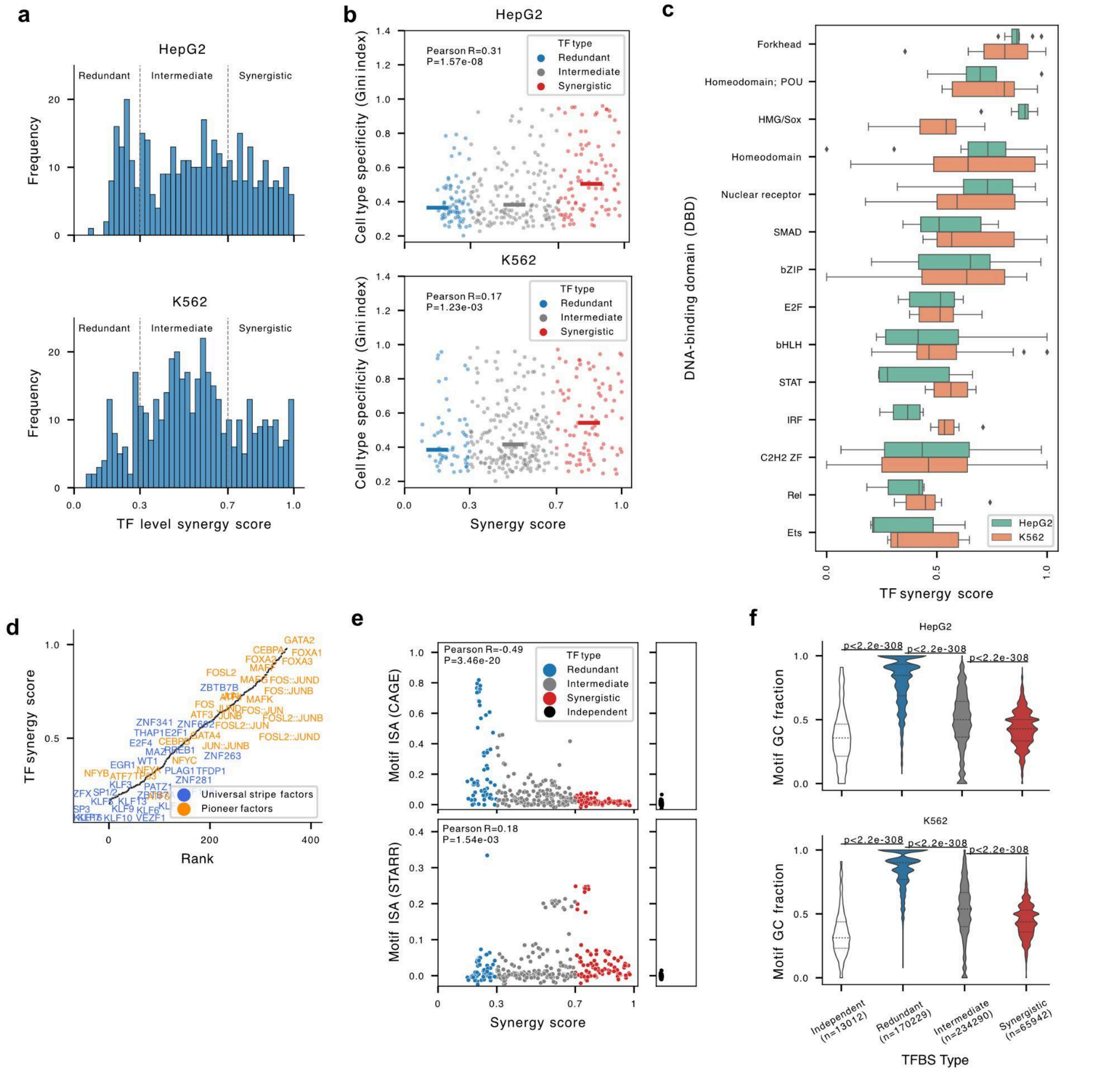


**Supplementary Figure 9. Properties of TF level synergy scores. a:** Histogram of TF-level synergy scores for HepG2 (upper ) and K562 (lower). Gray lines delineate redundant (synergy score < 0.3), intermediate (0.3 < synergy score < 0.7), and synergistic (synergy score > 0.7) TFs. **b:** Scatter plot for synergy score (horizontal axis) and cell type specificity measured by Gini index (vertical axis) of each TF in HepG2 (upper) and K562 (lower). Short solid lines indicate median synergy score of each TF type. **c:** Distribution of TF-level synergy score, grouped by DNA binding domain, colored by cell type. **d:** Ranked TF synergy scores with annotations of universal stripe factors (blue) and pioneer factors (orange) for HepG2. **e:** Scatter plot of synergy scores (horizontal axis) and motif ISA scores (vertical axis) of each TF in HepG2. Upper: motif ISA scores measured by CAGE track. Lower: motif ISA scores measured by STARR-seq track. **f:** Distribution of TF motif GC fraction for HepG2 (upper) and K562 (lower), grouped by TF type.

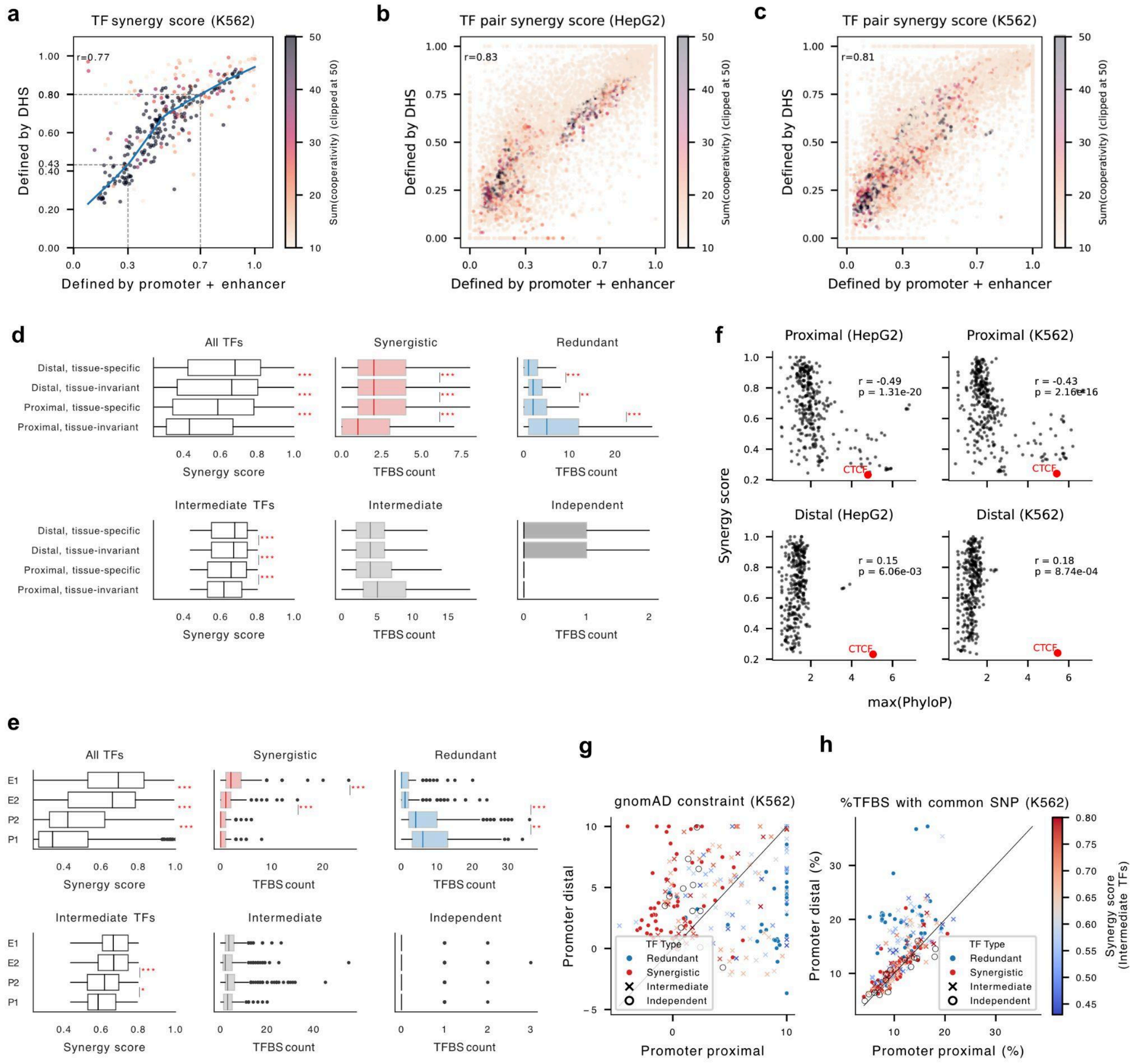


**Supplementary Figure 10. Differential contextual importance of redundancy and synergy. a:** Correlation between TF-level synergy scores defined by strict promoter and enhancer sets (horizontal axis, see Methods), versus synergy scores defined by DNase hypersensitive sites (DHS, vertical axis) for K562. **b-c:** Correlation between TF-pair-level synergy scores defined by strict promoter and enhancer sets (horizontal axis), versus synergy scores defined by DHSs (vertical axis), for HepG2 (**b**) and K562 (**c**). **d:** Left panel: distribution of TF-level synergy scores of all TFs (top), and intermediate TFs (bottom), grouped by region type. Middle and right panels: distributions of the number of binding sites per regulatory element for each TF type, grouped by region type. All measurements come from K562**.** Box plot: the box represents the interquartile range (IQR), with the median shown as a line inside the box. Whiskers extend to 1.5 times the IQR, and outliers beyond this range are not shown. ***: P≤1e-3, **: 1e-3≤P<1e-2, *:1e-2≤P<0.05. **e:** similar as **d**, but showing the distribution of synergy score and TFBS count by TF class for MPRA-defined E1, E2, P1, and P2. Dots represent outliers beyond 1.5x IQR. **f:** Scatter plot of max PhyloP scores versus TFBS (horizontal axis) and TF level synergy scores. **g:** gnomAD constraints for binding sites of each TF, calculated from promoter-proximal (horizontal axis) or promoter-distal (vertical axis) regions in K562. **h:** similar as d, but comparing the percentage of TFBS harboring at least one common SNP in promoter-proximal (horizontal axis) or promoter-distal (vertical) regions in K562. Redundant and synergistic TFs are shown as blue and red dots, respectively. Intermediate TFs are colored by their synergy score.

**Supplementary Table 1: ENCODE accession numbers for external datasets used in this study.**
https://github.com/anderssonlab/DeepCompARE/blob/main/Filter_regions/encode_accession.txt

**Supplementary Table 2: TF pair synergy scores for HepG2.**
https://github.com/anderssonlab/DeepCompARE/blob/main/Pd6_TF_cooperativity/tf_pair_synergy_score_hepg2_pe.csv

For column descriptions, see:
https://github.com/anderssonlab/DeepCompARE/blob/main/Pd6_TF_cooperativity/table_column_explanations.txt

**Supplementary Table 3: TF pair synergy scores for K562.**

https://github.com/anderssonlab/DeepCompARE/blob/main/Pd6_TF_cooperativity/tf_pair_synergy_score_k562_pe.csv

For column descriptions, see:
https://github.com/anderssonlab/DeepCompARE/blob/main/Pd6_TF_cooperativity/table_column_explanations.txt

**Supplementary Table 4: TF-level synergy scores for HepG2.**

https://github.com/anderssonlab/DeepCompARE/blob/main/Pd6_TF_cooperativity/tf_synergy_score_hepg2_pe.csv

For column descriptions, see:
https://github.com/anderssonlab/DeepCompARE/blob/main/Pd6_TF_cooperativity/table_column_explanations.txt

**Supplementary Table 5: TF-level synergy scores for K562.**

https://github.com/anderssonlab/DeepCompARE/blob/main/Pd6_TF_cooperativity/tf_synergy_score_k562_pe.csv

For column descriptions, see:
https://github.com/anderssonlab/DeepCompARE/blob/main/Pd6_TF_cooperativity/table_column_explanations.txt
